# Lizard feeding enhances *Ixodes pacificus* vector competency

**DOI:** 10.1101/2021.04.28.441694

**Authors:** Kacie Ring, Lisa Couper, Anne L. Sapiro, Fauna Yarza, X. Frank Yang, Keith Clay, Chase Mateusiak, Seemay Chou, Andrea Swei

## Abstract

A vector’s susceptibility and ability to transmit a pathogen— termed vector competency—determines disease outcomes, yet the ecological factors influencing tick vector competency remain largely unknown. *Ixodes pacificus*, the vector of *Borrelia burgdorferi* (*Bb*) in the western U.S., feeds on rodents, birds, and lizards. While rodents and birds are reservoirs for *Bb* and infect juvenile ticks, lizards are *Bb-*refractory. Despite *I. pacificus* feeding on a range of hosts, it is undetermined how larval host bloodmeal identity affects future nymphal vector competency. We experimentally evaluate the influence of larval host bloodmeal on *Bb* acquisition by nymphal *I. pacificus*. Larval *I. pacificus* were fed on either lizards or mice and after molting, nymphs were fed on *Bb-*infected mice. We found that lizard-fed larvae were significantly more likely to become infected with *Bb* during their next bloodmeal than mouse-fed larvae. We also conducted the first RNA-seq analysis on whole-bodied *I. pacificus* and found significant upregulation of antioxidants and antimicrobial peptides in the lizard-fed group. Our results indicate that the lizard bloodmeal significantly alters vector competency and gene regulation in ticks, highlighting the importance of host bloodmeal identity in disease transmission and upends prior notions about the role of lizards in Lyme disease transmission.

## Introduction

Vector competency—the ability of a vector to successfully acquire and transmit a pathogen—and the factors that modulate it are increasingly the focus of efforts to control the emergence and spread of vector-borne zoonotic diseases [1–5]. Manipulation of vector competency has been discussed as a disease prevention strategy in mosquitoes, teste flies, and triatome bugs [2, 3, 6]. In these vectors, rearing of naturally resistant populations, modifications of vector endosymbionts, and gene editing have been studied and implemented as applications of biological control to alter vector competency and prevent disease transmission. While strides have been made in understanding and manipulating vector competency in many systems, these studies highlight the complexity of vector-pathogen interactions and suggest that a more mechanistic understanding of disease transmission holds promise for disease control. For tick-borne pathogen systems in particular, the plasticity of vector competency and responsiveness to environmental or biological inputs remains poorly understood.

Tick-borne diseases constitute 40% of the emerging vector-borne diseases worldwide [7–9] and are sensitive to changing abiotic and biotic interactions driven by land use change and increased globalization [4, 8, 10, 11]. In the northern hemisphere, Lyme disease is the most common vector-borne disease, causing an estimated 300,000 cases annually in the U.S. [8, 12, 13]. It is caused by the bacterial agent *Borrelia burgdorferi* (*Bb*) and vectored by *Ixodes spp*. ticks, whose life history involves blood-feeding on a wide range of hosts during each of their three life stages (larvae, nymph and adult) [14]. Tick blood-feeding induces a suite of major physiological changes in the tick including antimicrobial activity [15] and cuticular reconstruction [16, 17]. In addition, the identity of the blood meal host has important consequences for pathogen acquisition, tick survivorship, and microbiome composition [18–21].

Mounting evidence indicates that microbiome composition impacts vector competency through induced immunological responses, morphological changes, or direct competition between microbial components of the tick microbiome [22–26]. However, the precise relationship between microbiome composition and tick competency for *Bb* is not well understood [27, 28]. Tick *Bb* acquisition is a complex process that requires the pathogen to evade numerous tick immune pathways and antimicrobial peptides [4, 6, 7, 8] followed by successful colonization of the midgut [9, 10]. There is evidence that these interactions may be influenced by biotic interactions such as host bloodmeal identity or microbiome interactions [11–13]. Manipulation of the microbiome in laboratory-reared *Ixodes scapularis* found that lower microbiome diversity reduced *Bb* colonization through induced changes in tick midgut morphology [24]. In *Ixodes pacificus*, greater microbiome diversity was associated with *Bb* colonization in one study but not another [19, 28]. The life history of *Ixodes pacificus*, the Lyme disease vector in the western U.S., provides a unique opportunity for natural microbiome manipulation. Juvenile *I. pacificus* feed predominantly on the western fence lizard, *Sceloporus occidentalis*, a *Bb*-refractory host, but will also parasitize reservoir competent hosts, typically rodents such as *Peromyscus* spp. mice, western gray squirrels (*Sciurus griseus*), and dusky-footed woodrats (*Neotoma fuscipes*) [29–31]. In addition to varying greatly in reservoir competency, blood meals from these host species can lead to stark differences in tick microbiome composition [19]. Lizard-feeding results in a significant reduction in *I. pacificus* microbiome diversity relative to rodent-feeding [19].

Given recent findings that lizard-feeding significantly reduces microbiome diversity [19] and experimental evidence of tick microbiome diversity affecting *Bb* colonization success [24], we sought to determine the direct effect of blood meal identity on *I. pacificus* vector competency in a tick pathogen acquisition experiment. We fed larval ticks on either lizards or mice then subsequently fed those ticks on *Bb*-infected mice and found that the ticks with previous lizard bloodmeals were significantly more susceptible to *Bb* infection. We then investigated mechanisms by which host blood meal may alter vector competency by conducting the first RNA-seq analysis on whole-bodied *I. pacificus* nymphs and comparing gene expression profiles for *I. pacificus* ticks following mouse or lizard larval blood meals. We find significant differences in tick vector competency based on larval host blood meal identity and detect multiple immune and metabolic factors that may alter *I. pacificus* vector competency for the Lyme disease pathogen.

## Materials and methods

### Ixodes pacificus collection

Fed *I. pacificus* larvae were collected from either western fence lizards (*Sceloporous occidentalis*) or deer mice (*Peromyscus maniculatus*). As lizards have naturally high larval burdens of *I. pacificus* (mean=25) [ref. 32], we collected ticks from *S. occidentalis* by capturing and holding lizards in drop off cages suspended over water for 3-4 days in a temporary field lab to collect replete ticks. We then transferred all collected, replete larvae to the lab facilities at San Francisco State University. Natural burdens of *I. pacificus* larvae on *Peromyscus spp*. are very low [32]. Because of low natural tick burdens, we experimentally attached up to 200 larval *I. pacificus* (BEI Resources, Manassas, VA) to *Peromyscus maniculatus* (Peromyscus Genetic Stock Center, Columbia, SC) in the lab. We collected and stored all replete ticks from lizard and mouse drop-off procedures under standard rearing conditions of 23°C and 90% relative humidity until they molted eight weeks later.

### Host inoculation and tick Bb acquisition experiments

*Borrelia burgdorferi* cultures were grown until they reached a concentration of 10^6^ spirochetes/mL [33]. Mice were inoculated intradermally with 100 μL of *Bb* culture (10,000 total spirochetes). Eight weeks post inoculation, successful *Bb* acquisition in the mice was determined via nested PCR of ear tissue targeting the 5S-23S rRNA spacer region [34].

Five C3H/HeJ mice were used to feed nymphs (Jackson Laboratory, Bar Harbor, Maine). Three of the C3H/HeJ mice were inoculated with *Bb* leaving the remaining two mice as uninfected controls. Nymphs that fed as larvae on either lizards or mice were then placed on either *Bb*-infected or uninfected C3H/HeJ mice for nymphal feeding. Host-to-tick acquisition experiments were conducted in three separate trials. The first trial was conducted at Indiana University, where lizard-fed and mouse-fed nymphs were fed on C3H/HeJ mice infected with *Bb* at a concentration of 10^5^ spirochetes/mL (1,000 total spirochetes). The second and third trials were conducted in the animal facilities at San Francisco State University where lizard-fed and mouse-fed ticks were subsequentially fed on C3H/HeJ mice infected with 10^7^ and 10^6^ spirochetes/mL (100,000 and 10,000 spirochetes total), respectively.

### Nucleic acid isolation and pathogen testing

After completing their bloodmeal, nymphs were placed under standard rearing conditions for 24 hours, flash frozen, and stored at −80°C until nucleic acid extraction. Prior to extraction, ticks were thoroughly surface sterilized with successive 1 mL washes of 3% hydrogen peroxide, 70% ethanol, and de-ionized H_2_O to remove surface contaminants. The tick was then lysed and homogenized using the Qiagen TissueLyser II (QIAGEN, Valencia, CA, USA). Tick samples were extracted simultaneously for total DNA and RNA using the Qiagen AllPrep DNA/RNA Micro Kit (QIAGEN, Valencia, CA, USA). DNA and RNA concentrations were measured using a Quibit Fluorometer (ThermoFisher Scientific, Walthman, CA, USA) in preparation for pathogen testing and library preparation. RNA content and quality were evaluated using a Bioanalyzer (Agilent, Santa Clara, CA, USA). DNA from engorged *Bb-*fed nymphs was tested for infection in triplicate by qPCR [35].

### Statistical analyses

To determine if larval bloodmeal host was a predictor of nymphal pathogen acquisition, we used a generalized mixed-effect model (GLMM) with a binomial error distribution. We used larval host bloodmeal (lizard or mouse feeding) as a fixed effect and trial as a random effect to account for experimental variation between trials. Analyses were performed using the glmm package (v.1.4.2) in R [36].

### Tick transcriptome analysis

Ticks from the third pathogen transmission experiment were used for transcriptome analysis. Unfed nymphs and engorged nymphs were divided into six experimental groups (Fig. 1). Our six experimental groups were composed of ticks that either fed on a lizard or a mouse as a larva. Then, molted nymphs from either larval bloodmeals either remained unfed, fed on an uninfected mouse, or fed on a *Bb-* inoculated mouse (Fig. 1). Each experimental group will hereinafter be described with the following abbreviations: unfed nymphs are referred to as “UF” and fed nymphs are referred to by whether they were fed on a *Bb*-positive “+Bb,” or *Bb*-negative “-Bb” C3H/HeJ mouse. Additionally, the larval bloodmeal (lizard or mouse) in each experimental group is indicated in the subscript following the abbreviation.

**Fig. 1.**
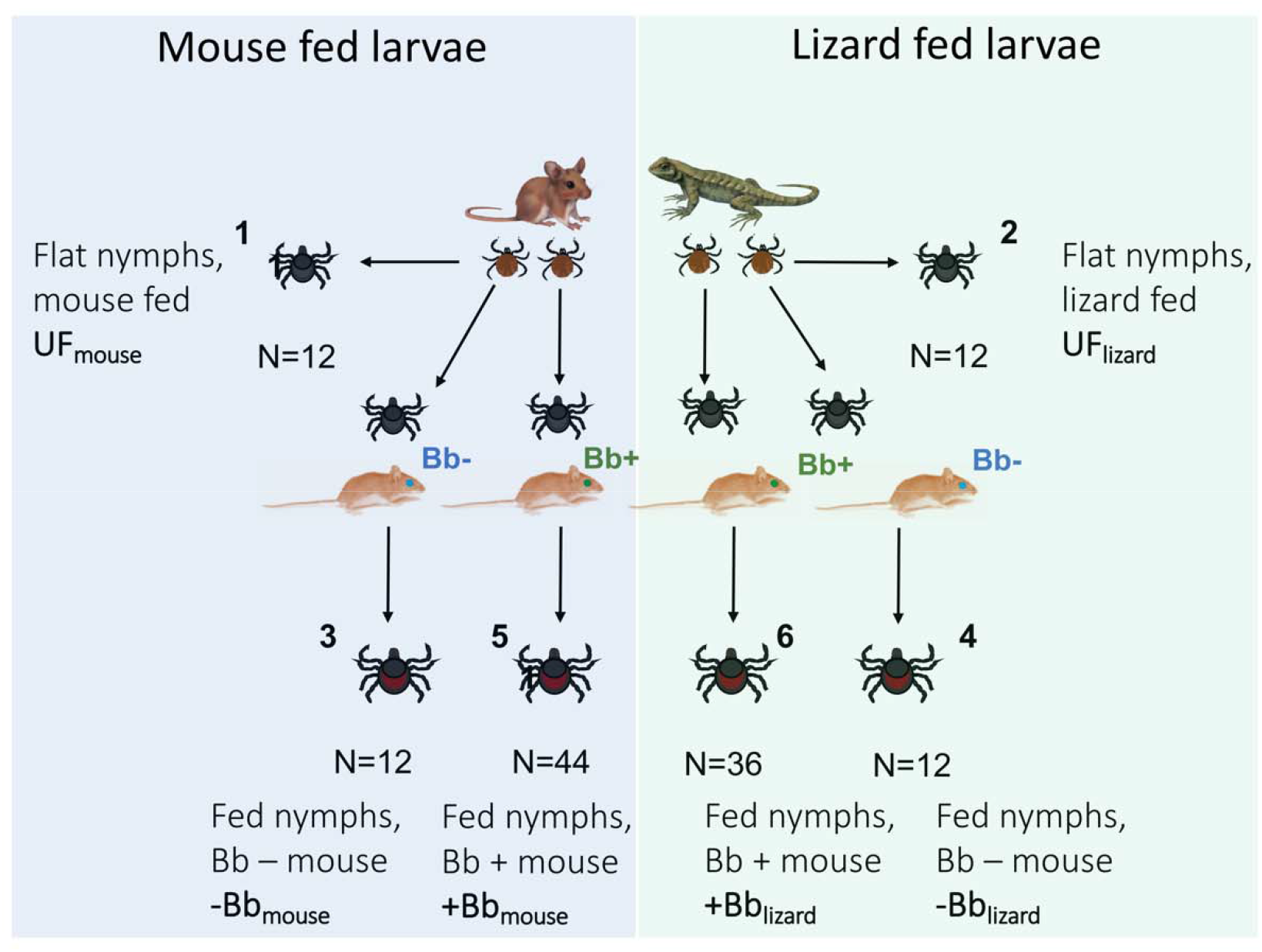
Transmission experiment design. Replete larval *I. pacificus* were obtained from *P. maniculatus* mice or *S. occidentalis* lizards. Successfully molted ticks from both groups were then either immediately sacrificed as unfed nymphs (groups 1&2) or fed on uninfected (groups 3 & 4) and *Bb*-infected (groups 5 & 6) C3H/HeJ mice. The *Bb-*fed ticks were analyzed via qPCR for *Bb* infection status. RNA from all groups was used to make RNA-seq libraries for transcriptomic analysis.

Groups one and two, which represent our unfed nymphs, “UF_lizard_” and “UF_mouse_” (Fig. 1), were set aside to examine the effect of the lizard or mouse larval bloodmeal on *I. pacificus* gene expression. The remaining four groups were engorged nymphal ticks. Uninfected control groups three and four, “-Bb_lizard_” and “-Bb_mouse_” were nymphs that fed on uninfected C3H/HeJ mice during their nymphal bloodmeal (Fig. 1). Groups five and six, “+Bb_lizard_” and “+Bb_mouse_” were nymphs that fed on *Bb* infected C3H/HeJ (Fig. 1).

We prepared three replicates from each of the six experimental tick feeding conditions (Fig. 1) for a total of 18 libraries. We pooled three individual ticks for each experimental replicate. RNA-seq libraries were prepared from total RNA extracted from ticks followed by rRNA depletion using Depletion of Abundant Sequences by Hybridization (DASH) [37]. The NEBNext Ultra II Directional RNA Library Prep Kit for Illumina (New England Biolabs, E7760S) was used to make RNA-seq libraries following the standard manual protocol. After constructing RNA-seq libraries from total RNA, reads containing rRNA sequences from *Ixodes spp*., *Bb*, and mouse were depleted using DASH, which targets Cas9 to abundant sequences in RNA-seq libraries. We utilized previously designed guide RNAs against mice and *Ixodes spp*. rRNAs [38], and we designed additional complementary guides to improve rRNA depletion of tick and *Bb* sequences using RNA-seq libraries made from total RNA from an *I. scapularis* nymph and *Bb* B-31 culture (Table S1).

### rRNA depletion with DASH

Guide RNAs were designed using DASHit (http://dashit.czbiohub.org/), and prepared from DNA oligos as in Gu et al. (2016) following the protocol for *In Vitro* Transcription for dgRNA Version 2 (dx.doi.org/10.17504/protocols.io.3bpgimn). The complete protocol for rRNA depletion with DASH can be found in the supplementary methods. Following DASH, RNA-seq libraries were sequenced on an Illumina NextSeq with paired-end 75 base pair reads. Fastq files and raw read counts have been deposited in the Gene Expression Omnibus (GEO) under accession number GSE173109.

### RNA-seq sequence processing and analysis

RNA-seq reads were trimmed of adapters and bases with quality score lower than 20 using Cutadapt [39] via Trim Galore! v0.6.5 and then mapped to the *Ixodes scapularis* ISE6 genome (assembly GCF_002892825.2_ISE6_asm2.2_deduplicated) accessed from NCBI RefSeq, using STAR (v2.7.3a) [40]. Reads mapping to predicted genes (gtf-version 2.2, genome build: ISE6_asm2.2_deduplicated, NCBI genome build accession: GCF_002892825.2, annotation source: NCBI Ixodes scapularis Annotation Release 100) were tabulated using Subread FeatureCounts (v2.0.0) [41], counting primary hits only. DESeq2 v1.26.0 [42], was used to determine differential expression between groups. Volcano plots from the ‘Enhanced Volcano” package in R, were used to visualize significant differential gene expression between experimental groups [43]. The visualization tools heat map and principle coordinate analysis plots in Deseq2 were utilized to visualize similarities across our gene expression profiles [42]. All code in our Deseq2 analysis is available at: https://github.com/choulabucsf/Ipac_DE_Ring_et_al_2021

## Results

### Host-to-tick Bb acquisition experiment

The effect of larval host bloodmeal on *I. pacificus* nymphal vector competency was examined in a host-to-tick pathogen acquisition experiment where replete larval *I. pacificus* were obtained from mice or lizards and then subsequently fed on *Bb*-infected C3H/HeJ mice (Fig. 1). A total of 36 lizard-fed (+Bb_lizard_) and 46 mouse-fed (+Bb_mouse_) *I. pacificus* nymphs were used across three experimental trials (Fig. S2).

*I. pacificus* nymphs that fed on *Bb*-inoculated C3H/HeJ mice were significantly more likely to become infected if they previously fed on lizards as larvae than if they fed on mice (χ^2^ (1, N = 82) = 7.8266, *p* =.0051). During their nymphal bloodmeal, 64% of lizard-fed ticks (N= 23/36) became infected with *Bb* compared to 30% of the mouse-fed ticks (N=14/46; Fig. 2). Even after accounting for trial as a random effect, our GLMM analyses found that the lizard larval bloodmeal is a significant, positive predictor of *Bb* acquisition in *I. pacificus* (Table 1). These results support our hypothesis that the host bloodmeal source may shape intrinsic vector competency of ticks across at least one life stage transition.

**Table 1.** Summary of results from a generalized mixed-effect model (GLMM) with binomial distribution examining the correlation between lizard larval blood host and the probability of infection in the nymphal stage. Inoculum load of host was included as a random effect to account for differences between experimental trials.

| Response | Fixed effect | Estimate | Standard Error | z value | p value |
| --- | --- | --- | --- | --- | --- |
| Infection status | (Intercept) | -1.02 | .51 | -2.023 | <.05* |
| Infection status | Lizard larval bloodmeal | 1.51 | .51 | 2.95 | <.01** |

**Fig. 2.**
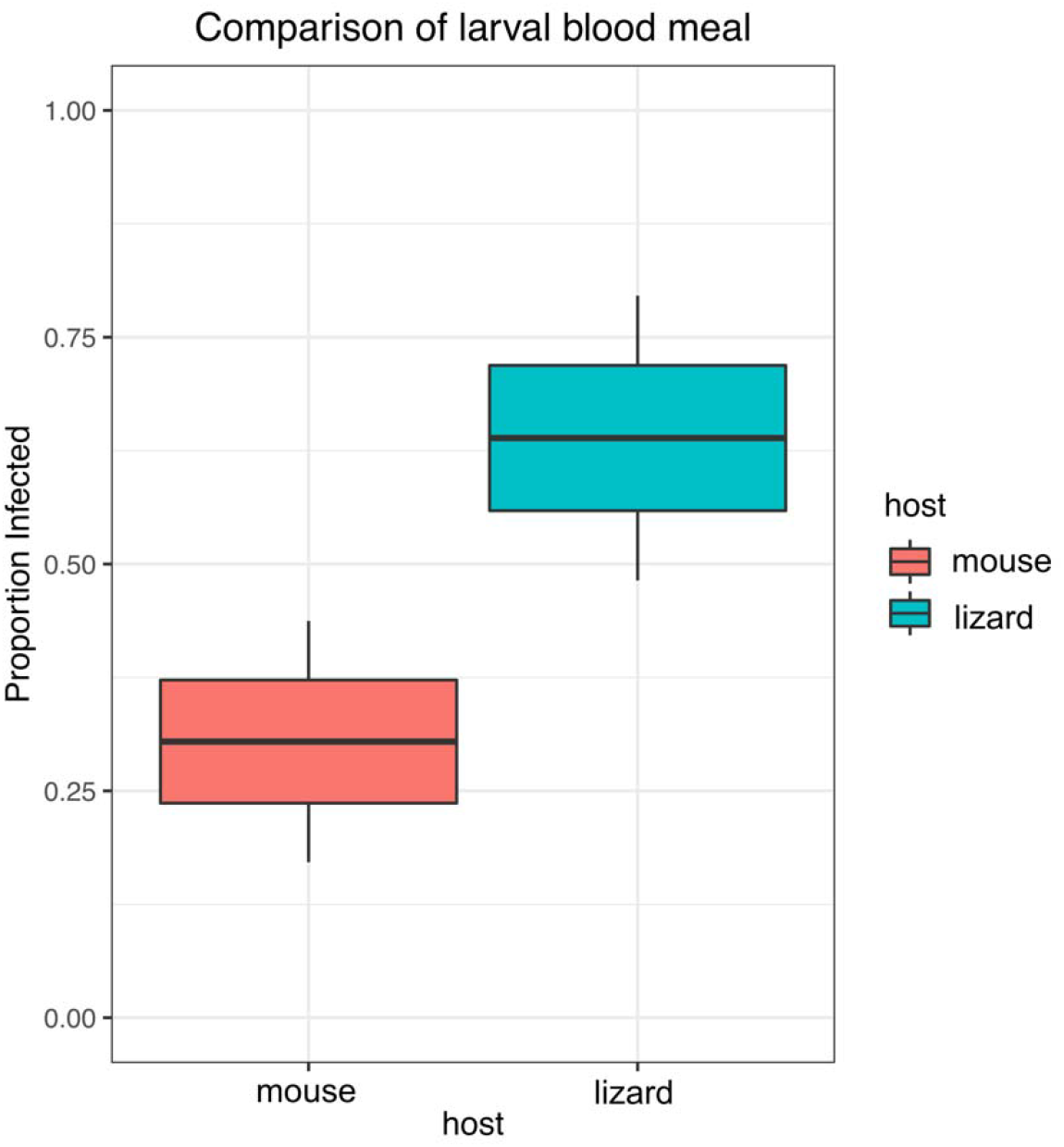
Comparison of infection status of *Bb*-fed *I. pacificus* nymphs with prior larval bloodmeals on either mice (mouse-fed; +Bb_mouse_) or lizards (lizard-fed; +Bb_lizard_). Lizard-fed larvae were significantly more likely to become infected when subsequently feeding on a *B. burgdorferi* infected mouse as nymphs than nymphs that previously fed on mice as larvae, with 64% of lizard-fed ticks infected compared to 30% of the mouse-fed ticks.

### Experimental transcriptomic differences

To investigate potential explanations for the differences in vector competency between lizard-fed and mouse-fed ticks, we conducted an RNA-seq analysis to compare gene expression between ticks with different blood histories and pathogen exposure. A total of 18 RNA-seq libraries were prepared from the six experimental groups, each represented by three replicates and resulting in over 370 million total reads (Fig. 1). With no available annotated *Ixodes pacificus* genome, we aligned our reads to the ISE6 *Ixodes scapularis* genome [44]. Mapping rates among replicates averaged at 45% with 15 million reads per library. Sequencing statistics for each replicate are presented in Table S2.

To visualize overall differences in gene expression profiles across the experimental groups, we created a heatmap and a PCA plot of our 18 replicates. The heatmap, generated from sample-to-sample distances, is based on read counts for all genes and showed that tick engorgement status (unfed vs. engorged) induced significant changes in *I. pacificus* gene expression (Fig. 3a). Additionally, the PCA plot indicated significant distinction of overall gene expression between unfed ticks of either bloodmeal type, UF_lizard_ and UF_mouse_ (Fig. 3b). The engorged experimental groups (groups three to six) had similar gene expression profiles and did not distinctly cluster together by experimental condition (Fig. 3b).

**Fig. 3.**
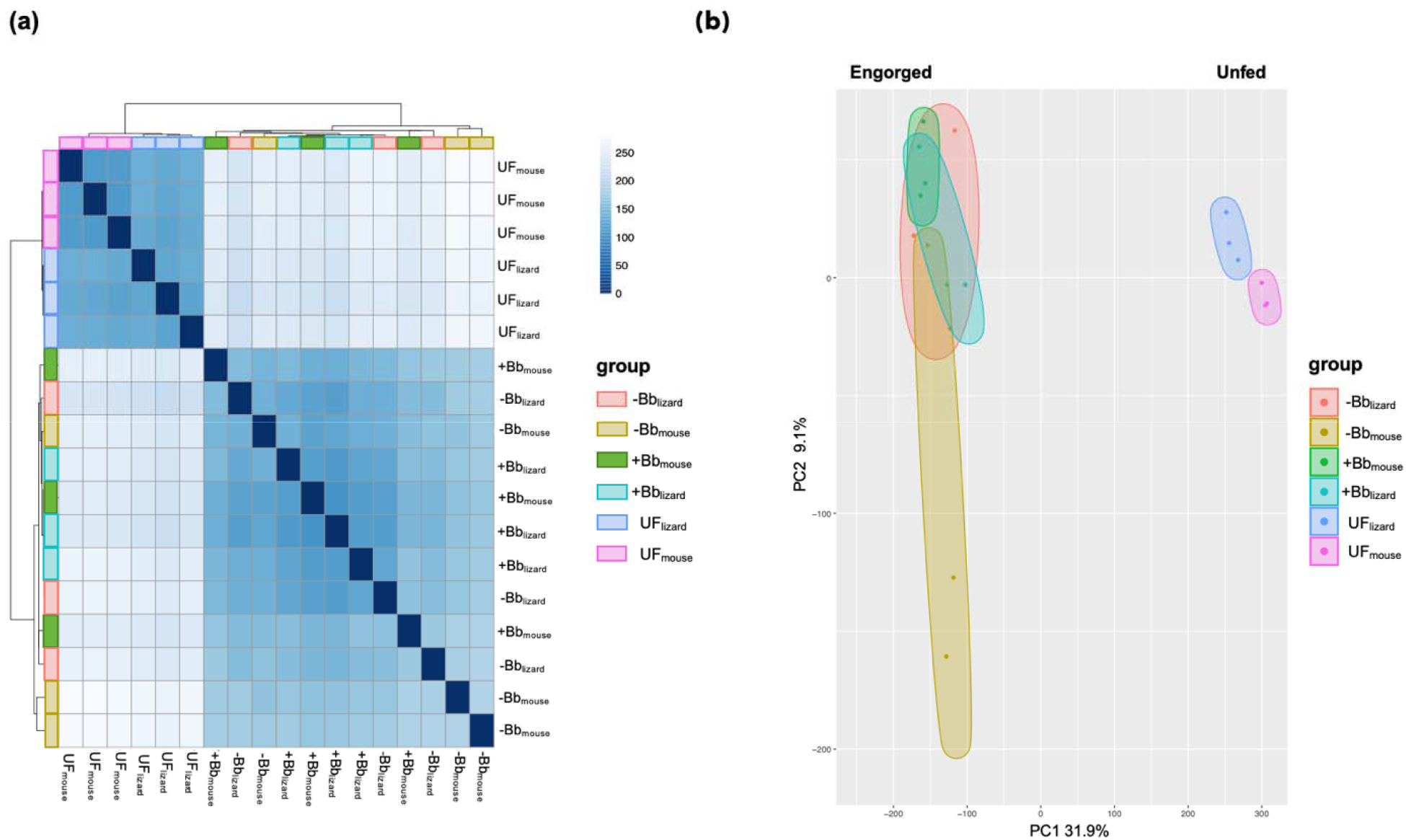
The similarity of transcriptomic profiles based on sample-to-sample distance shown by **(a)** a heat map plot of all samples. Visualization of the overall effect of experimental conditions shown by clustering in a **(b)** principal coordinate analysis on all transcriptomic profiles with plotted 95% confidence ellipses around experimental replicates.

### Differential gene expression

To investigate the mechanism through which host blood alters tick vector competency, we took a global transcriptomic approach to identify key genes or pathways modulated by mouse or lizard hosts. Differential gene expression analyses focused on several pairwise comparisons to examine transcriptomic differences between 1) the lizard versus mouse bloodmeal in the unfed group 2) unfed versus fed ticks, and 3) bloodmeal identity distinctions between *Bb* exposed groups.

The comparison between our unfed nymphs (UF_lizard_ vs. UF_mouse_), demonstrated that the lizard bloodmeal induced distinct transcriptomic changes in *I. pacificus* with 468 significantly differentially expressed genes (DEGs). While many of the DEGs remain undescribed, some of the highest upregulated genes induced by the lizard bloodmeal in the unfed group included antioxidants and antimicrobial peptides (Fig. 4a). The antioxidant glutathione peroxidase was the most significant DEG and was upregulated 48.5-fold after the lizard bloodmeal compared to mouse bloodmeal. Other tick antioxidants that were upregulated after the lizard bloodmeal include peroxidase (upregulated 21-fold) and glutathione-S-transferase (upregulated four-fold; Fig. 4a). We also found several DEGs that are related to the regulation of antimicrobial peptides but have never been described in *I. pacificus* ticks, such as acanthoscurrin-1, acanthoscurrin-2-like, micropulsin and micropulsin isoform, which were upregulated by 27.9, 104, 4, and 22.6-fold, respectively (Fig. 4a).

**Fig. 4.**
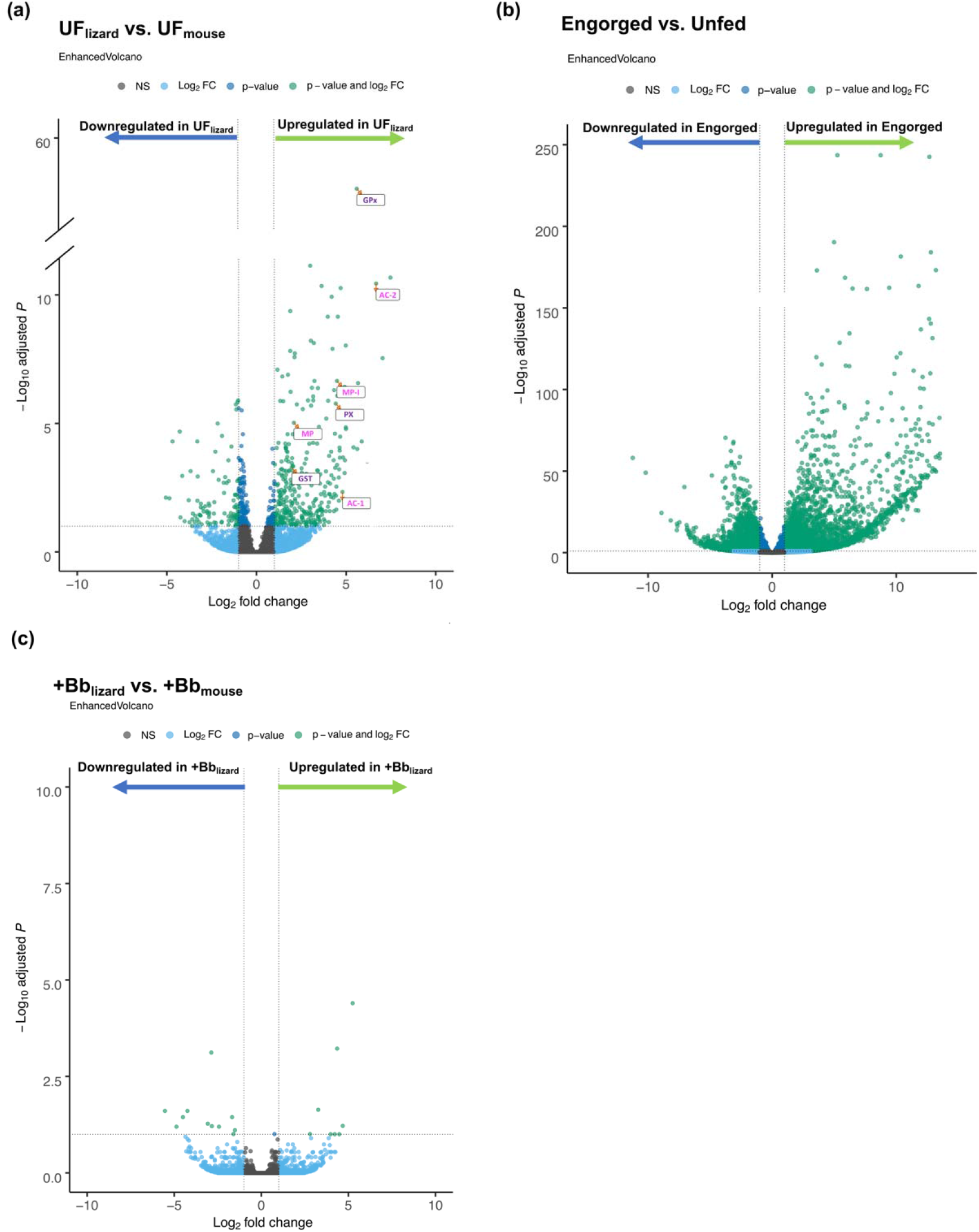
Volcano plots showing significant differential gene expression between the following: **(a)** UF_mouse_ (reference) vs. UF_lizard_ (comparison). Key annotated genes include tick antioxidants (purple print) glutathione peroxidase (GPx), glutathione S transferase (GST), peroxidase (PX) and antimicrobial peptides (pink print) acanthoscurrin-1 (AC-1), acanthoscurrin-2-like (AC-2), micropulsin (MP) and micropulsin isoform (MP-1). **(b)** Engorged (reference) vs. Unfed (comparison) and **(c)** +Bb_mouse_ (reference) vs. +Bb_lizard_ (comparison). Green points above the dotted x-axis represent genes significantly up or downregulated (padj value < 0.05 and log2FC > |1|).

To analyze the DEGs between engorged and un-engorged ticks, we combined the two unfed groups (UF_lizard_ & UF_mouse_) as our reference ‘unfed’ group and compared this group to all of the ‘engorged’ nymphs (i.e. −Bb_lizard_, −Bb_mouse_, +Bb_lizard_, & +Bb_mouse_). Engorgement induced significant gene expression differences, producing 6730 significant DEGs with a majority of difference in expression being upregulated genes in the engorged groups (Fig. 4b). Of the top 100 most significant DEGs, 25% were related to cuticle formation. Other notable genes that were differentially expressed include the antioxidant and detoxifying genes, glutathione peroxidase and sulfotransferase, which were both upregulated 4096-fold in the engorged group. Over a hundred DEGs between unfed and engorged ticks remain uncharacterized.

Despite significant differences in pathogen acquisition success between the +Bb_lizard_ and +Bb_mouse_ in pathogen transmission experiments, only 25 genes were differentially expressed between the two groups (Fig. 4c). The two most significantly DEGs included exonuclease V-like (upregulated 32-fold) and 4-coumarate –CoA ligase (downregulated 8-fold) in +Bb_lizard_ (Fig. 4c). No genes that are known to be related to immune function were detected as differentially expressed in the +Bb_lizard_ vs. +Bb_mouse_ comparison, with 7 of the 25 differentially regulated genes classified as uncharacterized.

## Discussion

Vector competency is considered an intrinsic property of a vector that determines its ability to acquire, maintain, and transmit pathogens [45], but the extent to which it is modulated by biotic or abiotic factors is poorly understood, especially in tick-borne pathogen systems. Here, we conducted a tick *Bb* acquisition experiment and transcriptome analysis on *I. pacificus* to determine if and by what potential mechanisms host bloodmeal history affects *I. pacificus* vector competency for the Lyme disease pathogen, *Bb*. Through *Bb* feeding experiments, we found that larval bloodmeal history significantly affects *I. pacificus* pathogen acquisition, a key component of vector competency. When ticks that fed on either lizards or mice as larvae fed on *Bb*-infected mice as nymphs, the previously lizard-fed ticks were twice as likely to acquire the pathogen. Further, significant transcriptomic signatures were detected between ticks with different bloodmeal histories. Gene expression analysis identified an upregulation of tick antioxidants and antimicrobial peptides in *I. pacificus* that fed on lizards, which may play a role in altering tick vector competency for *Bb*. Our results open the door for a potential mechanistic understanding of how host blood meal affects *I. pacificus* gene expression and the ecological factors that control *I. pacificus* susceptibility to *Bb*.

Recent studies suggest that tick microbiome composition can impact vector competency [17] and that host bloodmeal source can shape microbiome community structure [18, 23]. These two recent findings motivated this study to test whether the lizard bloodmeal host that has been previously shown to reduce tick microbiome diversity [19] can have subsequent effects on vector competency. Ticks with prior lizard or mouse bloodmeal histories displayed significant differences in pathogen acquisition when fed on mice infected with *Bb*. Across three separate experimental trials, we found that a prior lizard bloodmeal significantly increased the acquisition of *Bb* in nymphal *I. pacificus*. These results were surprising, especially given that infected *I. pacificus* that feed on *S. occidentalis* are cleared of their infection [46, 47]. The *Bb-*refractory nature of *S. occidentalis* has long been held as evidence of the lizard’s importance in maintaining lower prevalence of Lyme disease in the western U.S. and it likely contributes to lower disease risk relative to the northeastern U.S. However, whether this *Bb*-refractory property could be sustained transstadially in *I*. pacificus was unknown. Our results indicate that lizard-feeding does not preclude *Bb* infection in future life stages of *I. pacificus*, but rather enhances pathogen acquisition success relative to ticks with a prior mouse bloodmeal (Table 1). These results indicate that the acute and long-term consequences of a lizard bloodmeal on pathogen transmission are divergent.

The role of the microbiome in tick vector competency is unresolved [24, 28]. In a prior study, lower microbiome diversity in *I. scapularis* was associated with lower *Bb* colonization success due to decreased expression of genes involved in gut epithelium renewal, which enhances *Bb* colonization [24]. Therefore, we predicted that lizard-fed ticks, previously shown to have significantly lower microbiome species diversity than mouse-fed ticks [19], would similarly have lower *Bb* infection prevalence. However, our *Bb* acquisition experiment found that *I. pacificus* microbiome diversity, resulting from lizard-feeding [48], and pathogen transmission success are negatively correlated. This finding may be due to species-specific differences between *I. scapularis* and *I. pacificus* or be driven by the use of different experimental procedures used to manipulate the vector microbiome. Additionally, lizard feeding may affect tick vector competency through altering specific microbes rather than altering overall microbial diversity. Ultimately, the role of microbiome diversity and composition on pathogen acquisition success in *Ixodes* spp. remains uncertain and future studies are needed to disentangle the complicated interactions of these microbes.

Our RNA-seq analysis of *I. pacificus* with different bloodmeal histories revealed potential mechanisms that could be driving the differences observed in tick pathogen acquisition. Larval bloodmeal identity and engorgement have large impacts on *I. pacificus* gene expression (Fig. 3). Unfed nymphs clustered significantly by larval bloodmeal type (lizard vs. mouse; Fig. 3b), indicating that larval bloodmeal source induced distinct transcriptomic alterations in *I. pacificus*. We analyzed gene expression profiles in unfed nymphs right after they molted from larvae to nymph. Our analysis suggests that the effect of the larval bloodmeal on *I. pacificus* gene expression is carried through the transstadial molt and is present prior to the initiation of the nymphal bloodmeal. Among the unfed ticks, bloodmeal history drove divergence of 468 significantly expressed genes between un-engorged lizard and mouse fed ticks. The most significant DEG between unfed ticks with different bloodmeal histories was glutathione peroxidase in the lizard-fed group (Fig. 5a). Glutathione peroxidase is an important anti-oxidative enzyme, that works by reducing H_2_O_2_ and detoxifying OH radicals and prevents oxidative stress and cell damage in the tick [48, 49]. Two other known anti-oxidative enzymes, peroxidase and glutathione S-transferase were significantly upregulated after the lizard bloodmeal (Fig. 5b&c). A nutritional dependence on blood has required ticks to evolve and produce anti-oxidants to digest an inherently toxic meal containing high levels of iron and pro-oxidant levels [49]. Notably, glutathione peroxidase is homologous to SALP25d, a tick antioxidant produced in the salivary glands that has been shown to promote the transmission of *Bb* from tick to host and protects *Bb* from harmful hydroxyl radicals *in vitro* [16]. The upregulation of glutathione peroxidase in lizard-fed ticks has the potential to directly benefit *Bb* colonization from host to tick during the nymphal bloodmeal by increasing antioxidant concentration and protecting *Bb* from the harmful oxidative components of blood.

**Fig. 5.**
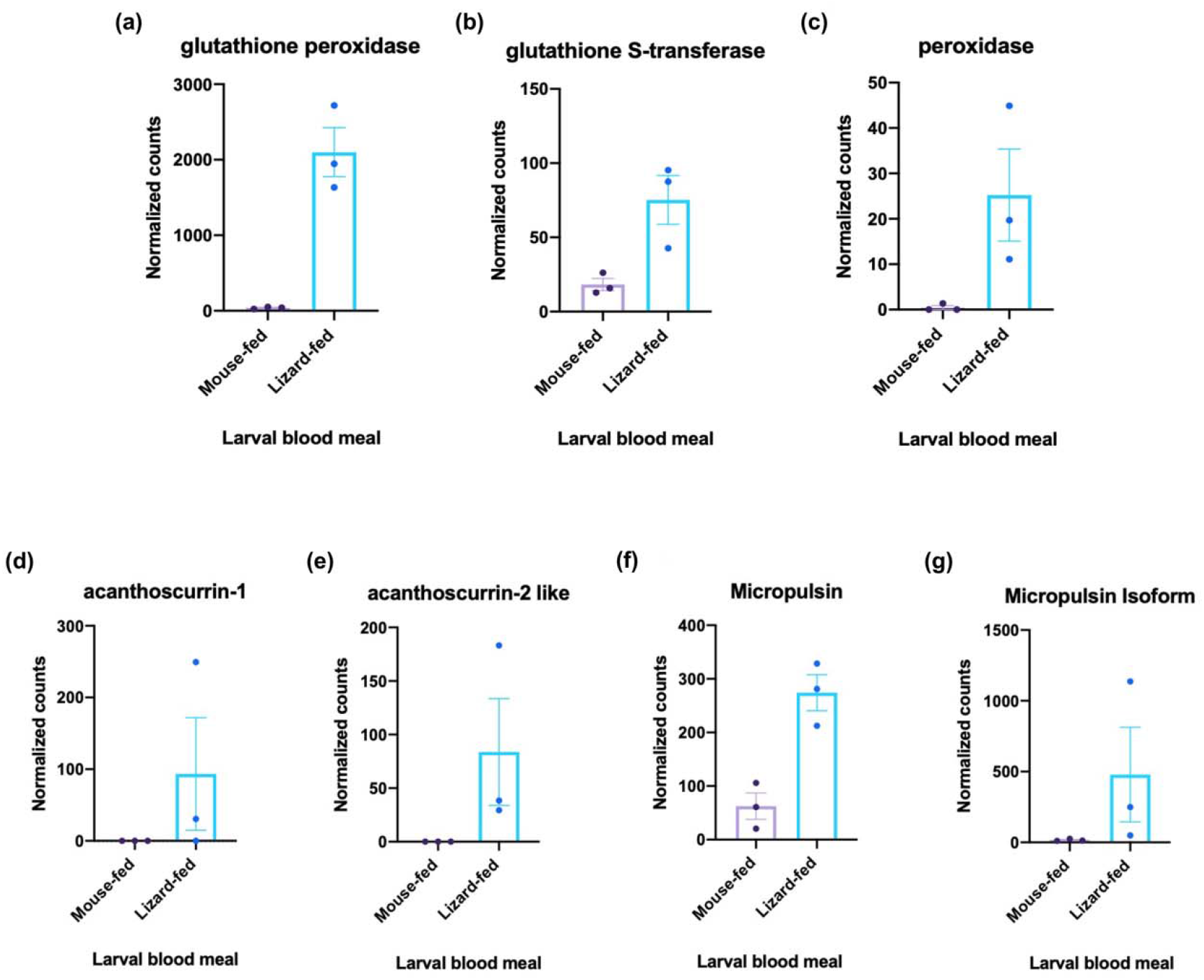
Comparison of key significant DEGs in unfed comparison UF_lizard_ vs. UF_mouse_. Graphs show the comparison of significantly DEGs using Wald’s test on normalized transcript for tick antioxidants **(a)** glutathione peroxidase (padj value= 2.79e-59) **(b)** glutathione-S-transferase (padj value=.0005) **(c)** peroxidase (padj value = 1.34e-6) and antimicrobial peptides **(d)** acanthoscurrin-1 (padj value=.003) **(e)** acanthoscurrin-2 like **(f)** micropulsin (padj value= 1.04e-5) **(g)** micropulsin isoform (padj value = 1.22e-7).

There was also a strong signal of microbial defense signals in unfed tick comparison. The antimicrobial peptides (AMPs) acanthoscurrin-1, acanthoscurrin-2, micropulsin, and a micropulsin isoform were all significantly upregulated in the unfed nymphs with prior lizard bloodmeals relative to prior mouse bloodmeals (UF_lizard_; Fig. 5d-g). Acanthoscurrin is a glycine-rich cationic AMP, known to be expressed in the hemocytes of tarantula spiders, *Acanthoscurria gomesiana*, and has activity against the yeast, *Candida albicans*, and gram-negative bacteria [51]. Micropulsin is a cysteine-rich AMP with histidine-rich regions, found in the hemolymph of the cattle tick, *Rhipicephalus microplus*, with high activity against gram-positive bacteria and fungus [52]. Neither of these AMPs have been detected in *Ixodes pacificus* prior to this study, but these results indicate that they may play an important role in pathogen acquisition and warrant further study. While lizard blood feeding contributes to the expression of AMPs, it is unclear what upstream components initiate their production. The antimicrobial activity may be an outcome of an initiated humoral immune response or derived from host immune effector molecules, as demonstrated when *Ixodes scapularis* feeds on a *Bb-*infected mouse [25]. To understand how the expression of these AMPs occur, future gene expression studies should examine the tick immune response to the lizard larval bloodmeal at multiple time points to track the immune response at different stages of feeding. Interestingly, the upregulation of AMPs with broad activity against microbes coincides with a previously described study showing that a lizard bloodmeal significantly reduces *I. pacificus* microbiome diversity after feeding [18]. Our results indicate that the lizard bloodmeal is associated with the production of AMPs that may reduce microbe-microbe competition for *Bb* colonization in future bloodmeals.

To further characterize the physiological changes that occur during *I. pacificus* host feeding, we analyzed unfed versus fed ticks, 24 hours after ticks completed their nymphal bloodmeal. Our study confirmed that engorgement induces a large number of transcriptional changes to the physical structure of the tick [46] (Fig. 3b). The greatest number of DEGs was between fed and unfed ticks (Fig. 4). Genes related to cuticle formation, antioxidant production, and detoxification were all significantly upregulated in fed ticks and are consistent with structural reformation that occurs during the engorgement process when ticks must rapidly synthesize a new cuticle over the course of taking a large bloodmeal [14]. Glutathione peroxidase and sulfotransferase were highly upregulated during engorgement and are critical for detoxifying the massive host bloodmeal and protect ticks from harmful oxidative stress inherent in blood feeding [47]. These results, while unsurprising, indicate that transcriptomic changes during *I. pacificus* engorgement are similar to the physiological alterations found in *Ixodes scapularis* [46].

Gene expression of *I. pacificus* is heavily shaped by engorgement status and bloodmeal history in unfed ticks but among the engorged nymphs (groups 5&6; Fig. 1), there was not a strong signal of bloodmeal history or infection status (Fig. 3b). Despite the significant differences in pathogen acquisition between host bloodmeal experimental groups (Fig. 2), only 25 genes with no known pathogen or immune function were differentially expressed between these groups. Comparing these results to our gene expression analysis from unfed nymphs, the strongest divergence in gene expression is present in the unfed ticks. This suggests that the physiological changes induced by the larval bloodmeal has lasting effects into the nymphal stage.

We document a strong correlation between host bloodmeal and vector competency, but there were limitations to our study. Naturally low burdens prevented us from using field-collected ticks for the mouse bloodmeal [32] and use of field-collected questing larvae is problematic because it is difficult to verify whether a tick had a previous, incomplete bloodmeal [27]. Our transcriptomic results indicate that bloodmeal history was only significantly different in the unfed nymphal group while fed nymphs were less apparent, which strongly suggests that larval host blood meal identity played a larger role in gene expression than tick source (Fig. 3).

Despite these intriguing results, our study highlights the importance of a more complete annotation of the reference transcriptome for *Ixodes spp*. ticks. A large proportion of DEGs remain uncharacterized indicating additional investigation into tick molecular function and transcriptomics is needed. The lack of differentially expressed genes in our comparison *of Bb*-exposed nymphs with different bloodmeal histories could be attributed to the timing of RNA sampling (24 hours after completed bloodmeal). Examination of gene expression before, during, and immediately after feeding would improve insight into the mechanism of pathogen colonization into *I. pacificus*. Additionally, future experiments should focus on understanding the role of antioxidants and the AMPs identified in this study in modifying tick vector competency. We found a strong association between lizard bloodmeal history and antioxidant activity as well as AMP production. These responses coupled with naturally high natural tick burdens and preferential feeding on lizards may suggest an evolutionary benefit to feeding on the western fence lizard for *I. pacificus* and perhaps *Bb*. Additional research should investigate whether changes in vector competency are propagated through additional life stages (i.e., nymphal to adult and adult to eggs), and its effect on the epigenetic memory of ticks over time [50].

The public health burden of Lyme disease is increasing, and diagnosis and treatment are expensive and imperfect. The complexity of tick-host-pathogen interactions involve many competing interactions, making intervention or prevention of disease transmission very difficult. A better understanding of the molecules, microbes, and antigens involved in vector competency presents a different approach to prevention [2, 3, 49, 51]. Identifying molecular and microbial drivers of tick survival and vector competency are attractive targets for novel control methods [4]. Using the perturbation of natural host bloodmeal, our study identified multiple molecular components that may be important in the successful acquisition of *Bb* in *I. pacificus* and identifies potential new targets for manipulating and preventing the transmission of tick-borne diseases.

## Acknowledgements

We thank the Chan Zuckerberg Biohub for sequencing. KR and AS want to acknowledge funding support from the Pacific Southwest Regional Center of Excellence for Vector-Borne Diseases funded by the U.S. Centers for Disease Control and Prevention (Cooperative Agreement 1U01CK000516. This work was also funded by NSF (#1427772, 1745411, 174037 to AS) and NIH (R01AI132851 to SC). Additional support for SC was provided by Chan Zuckerberg Biohub and the Pew Biomedical Scholars Program. FY is supported by the HHMI Gilliam Fellowship and ALS by the Life Sciences Research Foundation.

