## Supplemental information for "Lizard feeding enhances *Ixodes pacificus* vector competency"

Supporting Information
For

### Lizard feeding enhances *Ixodes pacificus* vector competency

Kacie Ring[1], Lisa Couper[2], Anne L. Sapiro[3], Fauna Yarza[3], X. Frank Yang[4], Keith Clay[5],

Chase Mateusiak[6], Seemay Chou[3, 6*], Andrea Swei[7*]

Institutional information:

1. Department of Ecology, Evolution, and Marine Biology, University of California, Santa Barbara, California, 93106
2. Department of Biology, Stanford University; 327 Campus Drive, Stanford, CA 94305
3. Department of Biochemistry and Biophysics, University of California, San Francisco; 600 16th Street, San Francisco, CA, 94158
4. Department of Microbiology and Immunology, Indiana University School of Medicine, 635, Barnhill Drive, MS409J, Indianapolis, IN 46202
5. Department of Ecology and Evolutionary Biology, Tulane University, 6823 St,. Charles Avenue, New Orleans, LA 70118
6. Center for Genome Science and Systems Biology, 4515 McKinley Ave, St. Louis, MO 63110
7. Chan Zuckerberg Biohub, San Francisco, CA, 94158
8. Department of Biology, San Francisco State University; 1600 Holloway Ave, San Francisco, CA 94132

#### Supplement Material

This file includes:

- Supplementary methods
- Figure S1
- Table S1
- Table S2

#### Supplementary methods

##### Experiments involving animals

All experiments involving *Peromyscus maniculatus* and C3H/HeJ mice at San Francisco State University were pre-approved by Institutional Animal Care and Use Committee (IACUC) under the protocol number AU19-01 and researchers were properly trained by the university veterinarian.

##### rRNA depletion with DASH

Both tracrRNA (5-AAAAAGCACCGACTCGGTGCCACTTTTTCAAGTTGATAACGGACTAGCCTTATTTTAACTTGCTATGCTGTCCTATAGTGAGTCGTATTA) and pooled crRNA DNA oligo templates (Table S1) were annealed to equimolar amounts of T7 primer (5’-TAATACGACTCACTATAG) by heating to 95ºC for 2 minutes and slowly cooling to room temperature. Annealed templates were used in 1 mL *in vitro* transcription reactions with: 120 µL 10X T7 buffer (400 mM Tris pH 7.9, 200 mM MgCl2, 50 mM DTT, 20 mM spermidine (Sigma 85558)), 100 µL of T7 enzyme (custom prepped enzyme gifted from E. Crawford, diluted 1:100 in T7 buffer, final concentration: 100 µg/mL), 300 µL NTPs (25 mM each, Thermo Fisher Scientific, R0481), 4 µg of annealed crRNA template or 8 µg of annealed tracrRNA template, and water to 1 mL. *In vitro* transcription was performed for 2 hours at 37°C. Reactions were split in half and purified using Zymo RNA Clean & Concentrator-5 Kit (Zymo Research R1015). To form the dgRNA complex for DASH, crRNA and tracrRNAs were diluted to 80 µM, mixed in equimolar amounts and annealed by heating to 95ºC for 30 seconds and cooling slowly to room temperature.

After transcription of dgRNAs, we performed DASH Protocol Version 4 (dx.doi.org/10.17504/protocols.io.6rjhd4n). All of our RNA-seq libraries were pooled in equimolar amounts to a final concentration of 2.8 nM. Cas9 and transcribed dgRNAs were prepped by mixing: 2.5 µL 10X Cas9 buffer (500 mM Tris pH 8, 1M NaCl, 100 mM $MgCl_2$, 10 mM TCEP), 2.5 µl 40 µM Cas9 (gift from E. Crawford, prepared as in Gu et al. (2016)), and 5 µL of 40 µM transcribed dgRNAs. The mix was incubated at 37°C for 5 minutes before 10 µL of pooled libraries were added. The mixture was incubated at 37°C for 1 hour and then purified with Zymo DNA Clean & Concentrator-5 (Zymo Research D4014) following the PCR product protocol and eluting DNA into 10.5 µL of water. During cleanup, Cas9 was again mixed with buffer and dgRNAs as above and incubated at 37°C for 5 minutes. Following cleanup, eluted DNA was added to the second Cas9-dgRNA mix, and mixture was incubated at 37°C for 1 hour for a second time. One $\mu$L of Proteinase K (New England Biolabs, P8107S) was added, and the mixture was incubated at 50°C for 15 minutes. The libraries were then purified with 0.9X volume of sparQ PureMag Beads (QuantaBio 95196) following the standard protocol, eluting in 23 µL of water. Libraries were amplified in a BioRad CFX96 using the Kapa HiFi Real-Time Amplification Kit in a 50 µL reaction with 25 µL master mix, 21 µl of the DASHed library pool, and 2 µL of 25 µM mix of Illumina P5 (5'-AATGATACGGCGACCACCGAGATCT) and P7 (5'-CAAGCAGAAGACGGCATACGAGAT) primers. The qPCR program for amplification was as follows: 98°C for 45 sec (1 cycle), [98°C for 15 sec, 63°C for 30 sec, 72°C for 45 sec, plate read, 72C for 20 sec] for 13 cycles. The libraries were removed from cycling conditions before leaving the exponential phase of amplification then purified with 0.9X volume of sparQ PureMag Beads according to the standard protocol.

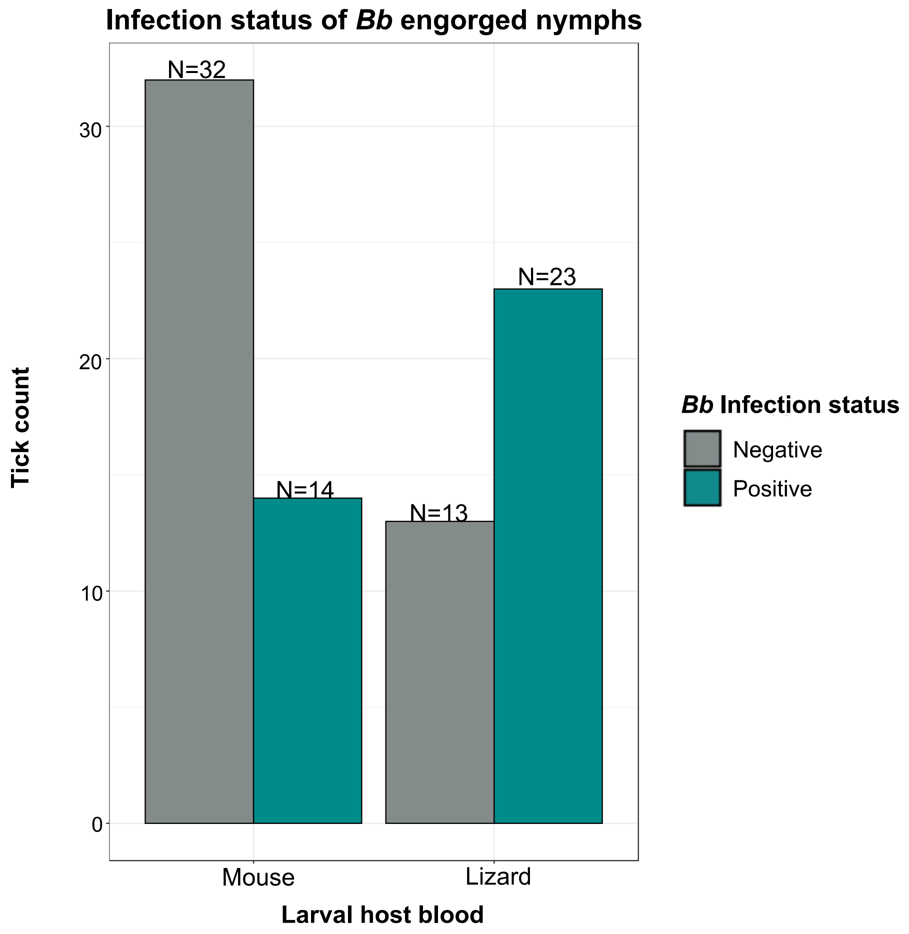


**Figure S1**. Comparison of infection status of *Bb*-fed *I. pacificus* nymphs with prior larval bloodmeals on either mice (mouse-fed; $+Bb_{mouse}$) or lizards (lizard-fed; $+Bb_{lizard}$). During their nymphal bloodmeal, 23 out of the 36 previously lizard-fed ticks became infected with *Bb* compared to 14 out of the 46 mouse-fed ticks.

**Table S1.** crRNA DNA oligo templates

| Guide Number | Target sequence | Reference |
|---|---|---|
| Tick_1 | GAGACTCTAGCCTATTAAAT | Dynerman et al. 2020 |
| Tick_2 | ACCATGGTTGTTACGGGTAA | Dynerman et al. 2020 |
| Tick_3 | AGGAGATCGAGTCCCTGGAA | Dynerman et al. 2020 |
| Tick_4 | GTGCTTGGTCGAGAGACCAG | Dynerman et al. 2020 |
| Tick_5 | GCCGCCGCTGGTGCAGATCT | Dynerman et al. 2020 |
| Tick_6 | TGGGTGGTAAACTCCATCTA | Dynerman et al. 2020 |
| Tick_7 | TCGGGGCTAGTTGCTTCGGC | Dynerman et al. 2020 |
| Tick_8 | GCAATTGTTCCCCTTGAACG | Dynerman et al. 2020 |
| Tick_9 | CGGGAAATGTGGTGTATGGG | Dynerman et al. 2020 |
| Tick_10 | AAAATTAGAGTGCTCAACGC | Dynerman et al. 2020 |
| Tick_11 | GCCAGAGGAAACTCTGGTGG | Dynerman et al. 2020 |
| Tick_12 | CACGTTGACATTCAGAGCAC | Dynerman et al. 2020 |
| Tick_13 | CTTTAAGAGTTTTAAGCAAG | Dynerman et al. 2020 |
| Tick_14 | ACCTATTCTCAAACTTTCAA | Dynerman et al. 2020 |
| Tick_15 | ACTCTGTGAAGAGACATGAG | Dynerman et al. 2020 |
| Tick_16 | TCTTCCTACTTGGATAACTG | Dynerman et al. 2020 |
| Tick_17 | GAGTAACTCAGGCGGATCGC | Dynerman et al. 2020 |
| Tick_18 | CTGCGGATGTGGGTACGGTC | Dynerman et al. 2020 |
| Tick_19 | GACGACTTAGGTACCTGTCG | Dynerman et al. 2020 |
| Tick_20 | CGCAGTAGTTCGTCTTGCGA | Dynerman et al. 2020 |
| Tick_21 | GTTCCGAGGACAGTGTCAGG | Dynerman et al. 2020 |
| Tick_22 | ACACTCTAAATCCTTTAACG | Dynerman et al. 2020 |
| Tick_23 | CTCCTTGTGGGGGCCCGTTC | Dynerman et al. 2020 |
| Tick_24 | GGGCGCCTTACCCCGGCGTT | Dynerman et al. 2020 |
| Tick_25 | CTGTGATGCCCTTAGATGTC | Dynerman et al. 2020 |
| Tick_26 | GTTTCCCGCTCCGCATCCGG | Dynerman et al. 2020 |
| Tick_27 | ATAACTTTGTGCTGATCGCA | Dynerman et al. 2020 |
| Tick_28 | GGAAGGTCCCGACGCTGGTC | Dynerman et al. 2020 |
| Tick_29 | TGCGATCCCGGGCCGCCCAG | Dynerman et al. 2020 |
| Tick_30 | GAAACCGCTTAGAGTAAAAC | Dynerman et al. 2020 |
| Tick_31 | AGATAGTAGATAGGGACAGT | Dynerman et al. 2020 |
| Tick_32 | TTCGGACCCCTGTGCGGTCC | Dynerman et al. 2020 |
| Tick_33 | CGTCGTCGAGCCGGGAATCG | Dynerman et al. 2020 |
| Tick_34 | AAGCAGAATTCGCCAAGCGT | Dynerman et al. 2020 |
| Tick_35 | GAAACCTCGCGTAGAGCAAA | Dynerman et al. 2020 |
| Tick_36 | CAAAACGGATCCGTAACCTT | Dynerman et al. 2020 |
| Tick_37 | AACTCAGAACTGGCACGGAC | Dynerman et al. 2020 |

| | | |
|---|---|---|
| Tick_38 | TCATCGTTTCTTTACTTACT | Dynerman et al. 2020 |
| Tick_39 | GGAGGGAACCAGCTACTAGA | Dynerman et al. 2020 |
| Tick_40 | GCGGTTAAAAAGCTCGTAGT | Dynerman et al. 2020 |
| Tick_41 | ATTCAATCGGTAGTAGCGAC | Dynerman et al. 2020 |
| Tick_42 | CGAGACTTGGTGTGAATTGC | Dynerman et al. 2020 |
| Tick_43 | CGGCGGGCTGCCGCGCTTCC | Dynerman et al. 2020 |
| Tick_44 | CCGCTGGAGGTGACCGGCGA | Dynerman et al. 2020 |
| Tick_45 | CCACATCCAAGGAAGGCAGC | Dynerman et al. 2020 |
| Tick_46 | ACGGGCCTGGCACCCACTCT | Dynerman et al. 2020 |
| Tick_47 | CGAACTTCTCGTGCACGGCG | Dynerman et al. 2020 |
| Tick_48 | GGCACGCAGACGCGGTCTCC | Dynerman et al. 2020 |
| Tick_49 | CGGGTTGGTCCCGTGGCGGC | Dynerman et al. 2020 |
| Tick_50 | GGTGCATGGCCGTTCTTAGT | Dynerman et al. 2020 |
| Tick_51 | GTTGCCGCACTGGGTCCCGA | Dynerman et al. 2020 |
| Tick_52 | GTGCATCTGCCGGCAGCGGC | Dynerman et al. 2020 |
| Tick_53 | AAAACTAACCTGTCTCACGA | Dynerman et al. 2020 |
| Tick_54 | GTTTCTGTTCTGTTGGTTTT | Dynerman et al. 2020 |
| Tick_55 | GGCCCTTGAAAATCCGAGTG | Dynerman et al. 2020 |
| Tick_56 | TTCAGTCGTAATCCCACGGA | Dynerman et al. 2020 |
| Tick_57 | GAGGCGGGTTCTTTCCGCGG | Dynerman et al. 2020 |
| Tick_58 | TGCCGGCGACGGCCGGGGAT | Dynerman et al. 2020 |
| Tick_59 | GCGGAGGAAAAGAAACCAAC | Dynerman et al. 2020 |
| Tick_60 | GAGAACCTTGAGGACTGAAG | Dynerman et al. 2020 |
| Tick_61 | CTGAAACTTAAAGGAATTGA | Dynerman et al. 2020 |
| Tick_62 | CTGATTTAATGAGCCATTCG | Dynerman et al. 2020 |
| Tick_63 | TAATTAGTGACGCGCATGAA | Dynerman et al. 2020 |
| Tick_64 | ACATCATGCCTGTCGTGGCT | Dynerman et al. 2020 |
| Tick_65 | TTCAGTCCGAGCCGTCGCAT | Dynerman et al. 2020 |
| Tick_66 | TCGCTCCTCCTTAGGCGACC | Dynerman et al. 2020 |
| Tick_67 | TGGTCCTTCGGGGCCCAGTG | Dynerman et al. 2020 |
| Tick_68 | GGCATAGTTCACCATCTTTC | Dynerman et al. 2020 |
| Tick_69 | AGACAGCAGGACGGTGGCCA | Dynerman et al. 2020 |
| Tick_70 | AAAGTTACCACAGGGATAAC | Dynerman et al. 2020 |
| Tick_71 | TTCTTTTCAACTTTCCCTCA | Dynerman et al. 2020 |
| Tick_72 | GGTAAGCAGAACTGGCGCTG | Dynerman et al. 2020 |
| Tick_73 | TAACATGTGCGCGAGTCAAT | Dynerman et al. 2020 |
| Tick_74 | TGCAGTGCCGTTCCTGCTTC | Dynerman et al. 2020 |
| Tick_75 | CTGGTCGCTCTCCGGGGCTA | Dynerman et al. 2020 |
| Tick_76 | CCGCACGCGCGCTACACTGA | Dynerman et al. 2020 |

| | | |
|---|---|---|
| Tick_77 | TGCTGCCTCCCGTAGGAGTC | Dynerman et al. 2020 |
| Tick_78 | TTTATTAGACCAAGATCAAT | Dynerman et al. 2020 |
| Tick_79 | ATTAAAACAAAGCATTGCGA | Dynerman et al. 2020 |
| Tick_80 | CGGTGAAATGCGCAGATATC | Dynerman et al. 2020 |
| Tick_81 | GACCAAACTTGATCATTTAG | Dynerman et al. 2020 |
| Tick_82 | TATTGGAGCTGGAATTACCG | Dynerman et al. 2020 |
| Tick_83 | TTGCTCTGTTTCGTGCGGCT | Dynerman et al. 2020 |
| Tick_84 | AAGCGAATGATTAGAGGCAT | Dynerman et al. 2020 |
| Tick_85 | TAAGTTCCGACCTGCACGAA | Dynerman et al. 2020 |
| Tick_86 | CCGCGCGGTGCCGCGCACTC | Dynerman et al. 2020 |
| Tick_87 | TTCACCTTGGAGACCTGCTG | Dynerman et al. 2020 |
| Tick_88 | CACACCAAAAACGAGTGCAT | Dynerman et al. 2020 |
| Tick_89 | AACGAAAGTTAGAGGTTCGA | Dynerman et al. 2020 |
| Tick_90 | AGCTTATGACTCGCGCTTAC | Dynerman et al. 2020 |
| Tick_91 | GGGGCGGTACATCTGTCAAA | Dynerman et al. 2020 |
| Tick_92 | CGCGGCGTTCTACCGGCGGT | Dynerman et al. 2020 |
| Tick_93 | GGCCGAAACCCGATCGATCT | Dynerman et al. 2020 |
| Tick_94 | TAAAACTCAAAGGAATTGAC | Dynerman et al. 2020 |
| Tick_95 | AGGGAGTAGCCCATGAGGCC | Dynerman et al. 2020 |
| Tick_96 | CGCTAAGCTGGGCGTCGATA | Dynerman et al. 2020 |
| Tick_97 | GGTGCAAGTCCCCCTGACAG | Dynerman et al. 2020 |
| Tick_98 | TTGAGAGCTCTTTCTTGATT | Dynerman et al. 2020 |
| Tick_99 | GACTAGAGTCAAGCTCAACA | Dynerman et al. 2020 |
| Tick_100 | CGGAGAGGGAGCCTGAGAAA | Dynerman et al. 2020 |
| Tick_101 | CGCTGCACCTAAATGCATTT | Dynerman et al. 2020 |
| Tick_102 | CTACGTATTACCGCGGCTGC | Dynerman et al. 2020 |
| Tick_103 | GCAGTTGAACATGGGTCAGT | Dynerman et al. 2020 |
| Tick_104 | GGGCTGTTCGCCCATTAAAG | Dynerman et al. 2020 |
| Tick_105 | GCAAAATGCCCCCGTAACTT | Dynerman et al. 2020 |
| Tick_106 | GCTGCGGTGAATACGTTCCC | Dynerman et al. 2020 |
| Tick_107 | GCGGCCTATCAGCTTGTTGG | Dynerman et al. 2020 |
| Tick_108 | TCGCAGGCTCATTCTTCAAA | Dynerman et al. 2020 |
| Tick_109 | CTTTGCAACCATACTTCCCC | Dynerman et al. 2020 |
| Tick_110 | GCCCAAAGGTTCCCTCAGCC | Dynerman et al. 2020 |
| Tick_111 | AACCGCTGAAAGCATCTAAG | Dynerman et al. 2020 |
| Tick_112 | GTGACGAAAAATAACAATAC | Dynerman et al. 2020 |
| Tick_113 | GAAGTTTCCCTCAGGATAGC | Dynerman et al. 2020 |
| Tick_114 | TGATCAAGAACGAAAGTCAG | Dynerman et al. 2020 |
| Tick_115 | ATCGCTTGGCGCTAGAAGCG | Dynerman et al. 2020 |

| | | |
|---|---|---|
| Tick_116 | GGGGTCAACTCGGAGGAAGG | Dynerman et al. 2020 |
| Tick_117 | AGAGTTTTCTTTTCTCTGTA | Dynerman et al. 2020 |
| Tick_118 | TAGACGGGATTCTGACTTAG | Dynerman et al. 2020 |
| Tick_119 | GCGAACAGGATTAGATACCC | Dynerman et al. 2020 |
| Tick_120 | TTGAACGCACATTGCGGCCT | Dynerman et al. 2020 |
| Tick_121 | GGCGGTGTGTACAAAGGGCA | Dynerman et al. 2020 |
| Tick_122 | TAATGATTAAGAGGGACAGA | Dynerman et al. 2020 |
| Tick_123 | AGTTGTTACACACTCCTTAG | Dynerman et al. 2020 |
| Tick_124 | CAGCTCGTGTCGTGAGATGT | Dynerman et al. 2020 |
| Tick_125 | CCTACCATCGAAAGTTGATA | Dynerman et al. 2020 |
| Tick_126 | ACGCCTGTCGGCCTCGCCTT | Dynerman et al. 2020 |
| Tick_127 | GTTCGGGCCTCCCGACAGCC | Dynerman et al. 2020 |
| Tick_128 | GCCCCACACGCTCACGTCGC | Dynerman et al. 2020 |
| Tick_129 | GGCATGTACTTAGACATGCA | Dynerman et al. 2020 |
| Tick_130 | GACTGACGCCTGCCCGGTGC | Dynerman et al. 2020 |
| Tick_131 | GATCCGAGGATGTCCGAATG | Dynerman et al. 2020 |
| Tick_132 | ATAGTTTGCGACCTCGATGT | Dynerman et al. 2020 |
| Tick_133 | GCTGCCGCCAGTGCAGATCT | Dynerman et al. 2020 |
| Tick_134 | AGATCAGCCTGTTATCCCCG | Dynerman et al. 2020 |
| Tick_135 | GGTGAAAAGTACCCCGGGAG | Dynerman et al. 2020 |
| Tick_136 | GCTTTTTGATCCTTCGATGT | Dynerman et al. 2020 |
| Tick_137 | AGAGCGCGGCAGTGAGACTG | Dynerman et al. 2020 |
| Tick_138 | GGTAAGCAGAACTGGCGATG | Dynerman et al. 2020 |
| Tick_139 | GCTAGAGTCTCGTTCGTTAT | Dynerman et al. 2020 |
| Tick_140 | CGGGTGGTAAACTCCATCTA | Dynerman et al. 2020 |
| Tick_141 | CTGTGTAGTTAGCTGGCAAT | Dynerman et al. 2020 |
| Tick_142 | CGAACGGGTGAGTAACACGT | Dynerman et al. 2020 |
| Tick_143 | GCGGTGAGCGTTGAAGGTGT | Dynerman et al. 2020 |
| Tick_144 | TGAAACATCTCAGTACCCGC | Dynerman et al. 2020 |
| Tick_145 | ACCACCAAGATCTGCACTAG | Dynerman et al. 2020 |
| Tick_146 | TGTGGAAAGCCTCAGCTGCG | Dynerman et al. 2020 |
| Tick_147 | TCGCCCCGAGATAGGGATGG | Dynerman et al. 2020 |
| Tick_148 | GGGCGTAAAGAGCTCGTAGG | Dynerman et al. 2020 |
| Tick_149 | AAACTATGCCGACTAGGGAT | Dynerman et al. 2020 |
| Tick_150 | GGACTATCGGCTCAAGCCGA | Dynerman et al. 2020 |
| Tick_151 | CTCTCAAGCAACCCGACTCA | Dynerman et al. 2020 |
| Tick_152 | GGGGGATTCGGTGCTGGGCT | Dynerman et al. 2020 |
| Tick_153 | CTGTGGCTGCCCCCGTCAAG | Dynerman et al. 2020 |
| Tick_154 | TCACTCTACTTGTGCGCTAT | Dynerman et al. 2020 |

| | | |
|---|---|---|
| Tick_155 | AAAATTAGAGTGTTCAAAGC | Dynerman et al. 2020 |
| Tick_156 | CTCCCGCGGAGGCGCTATCG | Dynerman et al. 2020 |
| Tick_157 | GGCGTCCGCAATGAATCTTG | Dynerman et al. 2020 |
| Tick_158 | GAGTAACTATGACTCTCTTG | Dynerman et al. 2020 |
| Tick_159 | GACGAGTGAGCGCCTGGGAC | Dynerman et al. 2020 |
| Tick_160 | TACTGGGAATTCCTCGTTCA | Dynerman et al. 2020 |
| Tick_161 | AGTGCCAAGTGGGCCACTTT | Dynerman et al. 2020 |
| Tick_162 | GATTCAGTCTTGGTGTGACG | Dynerman et al. 2020 |
| Tick_163 | AAGCCGGTCTCAGTTCGGAT | Dynerman et al. 2020 |
| Tick_164 | GGCACGAAGCTACCATCCGT | Dynerman et al. 2020 |
| Tick_165 | AGCGAAAGCGAGTCTGAATA | Dynerman et al. 2020 |
| Tick_166 | GTAGGACGCCGCGACGGGCA | Dynerman et al. 2020 |
| Tick_167 | AGCGGAGACTGCCATTTACG | Dynerman et al. 2020 |
| Tick_168 | GCACAGCTGTGGCTTTCCAC | Dynerman et al. 2020 |
| Tick_169 | CTCTACCTGACCACCTGTGT | Dynerman et al. 2020 |
| Tick_170 | AGAGTTTCTCGGGTCGATCG | Dynerman et al. 2020 |
| Tick_171 | CTCAGTACGAGAGGAACCGC | Dynerman et al. 2020 |
| Tick_172 | GTGTCGGGCGTGAGCCCGCC | Dynerman et al. 2020 |
| Tick_173 | GTGAATCTGCCGGGACCACC | Dynerman et al. 2020 |
| Tick_174 | AGAGGTGTAGCATAAGTGGG | Dynerman et al. 2020 |
| Tick_175 | GGAGTTAGCAAATGCACGAG | Dynerman et al. 2020 |
| Tick_176 | ATCGAGGGGCCCCGCTTTCG | Dynerman et al. 2020 |
| Tick_177 | GATCCGAGGGTGTCCGAATG | Dynerman et al. 2020 |
| Tick_178 | AAAGAAACGTCGCCGGCTCG | Dynerman et al. 2020 |
| Tick_179 | TGTCTTCGGGCGCAACAAGA | Dynerman et al. 2020 |
| Tick_180 | AAACGTCAGGGGGAAATACC | Dynerman et al. 2020 |
| Tick_181 | GAGGGATTCGGTGCTGGGCT | Dynerman et al. 2020 |
| Tick_182 | CAGAGGAATTTGCTCTGTAA | Dynerman et al. 2020 |
| Tick_183 | CGTAGCCTCGGGTCACCTTC | Dynerman et al. 2020 |
| Tick_184 | AAACAGCTTGCCTGGAGATA | Dynerman et al. 2020 |
| Tick_185 | TAGAGGTGAAATTCTTGGAC | Dynerman et al. 2020 |
| Tick_186 | CCAATACCAAGCTATAGTAA | Dynerman et al. 2020 |
| Tick_187 | TAATAAATGCGTCCGCCTTG | Dynerman et al. 2020 |
| Tick_188 | AAGAGTGCGTAATAGCTCAC | Dynerman et al. 2020 |
| Tick_189 | TAGAACGTCGTGAGACAGTT | Dynerman et al. 2020 |
| Tick_190 | ATTGTGTGCCTCGAGAAGGC | Dynerman et al. 2020 |
| Tick_191 | AGAATTACTATCGCAACGAC | Dynerman et al. 2020 |
| Tick_192 | TGGTCCTTCGGGGCCCAGTG | This work |
| Tick_193 | TCGCAACTCGGCACGGCACG | This work |

| | | |
|---|---|---|
| Tick_194 | CCCCACACGCTCACGTCGTG | This work |
| Tick_195 | TCAGGCGAATACCTGGCGTA | This work |
| Tick_196 | ATCGTTTCGGGCCTCCCCGA | This work |
| Tick_197 | ACTCCCACGACAGCCAAACA | This work |
| Tick_198 | CGCGGCACCGCGCGGGCAGC | This work |
| Tick_199 | AGTTTCGCTGTGACGTGGAG | This work |
| Tick_200 | ACACTCTAAATCCTTTAACG | This work |
| Tick_201 | GACGCCTTTCGTCGCCGTCT | This work |
| Tick_202 | CGGACACCCACCGGCTAAAT | This work |
| Tick_203 | TTTGCACGTCAGAAGCGCTT | This work |
| Tick_204 | GAACACCCTCAACCACGATG | This work |
| Tick_205 | CGCAATGAAACGTGAAGGCG | This work |
| Tick_206 | GTCTTGAAAACGAATGCGAG | This work |
| Tick_207 | GTCATCAATGGCAGGAACCC | This work |
| Tick_208 | ACTCAAAGTCGTTCTTGAAA | This work |
| Tick_209 | AGTGCGTTCTGACCGAAGGC | This work |
| Tick_210 | TAGAGGAATTTGCTCTGTGA | This work |
| Tick_211 | GCTCCCGCGTCCTGCAGTAG | This work |
| Tick_212 | ATTTACTTGTATAACCGCGG | This work |
| Tick_213 | ATCTCATGAGCGTCCCGCAT | This work |
| Tick_214 | ATCTCACTTCTGAGAGCGAC | This work |
| Tick_215 | TGCGGCCTCGGGCCTTCCCG | This work |
| Tick_216 | TTTCCCGCTCCGCATTCCGG | This work |
| Tick_217 | TTTGCGTTCAACTTTTCACC | This work |
| Tick_218 | ATTCAATATAATTAATATAG | This work |
| Tick_219 | GGCGTCGTCAATGATTTTTT | This work |
| Tick_220 | TGCACTTAGTGCTAGAAGCG | This work |
| Tick_221 | GTTCGCAGCTAGGCTTAGCG | This work |
| Tick_222 | GGCAGGGGTCAGCACGGCCG | This work |
| Tick_223 | TCTGGGTGCCACCGCAACAG | This work |
| Tick_224 | AGCCCGATGCTGCTTAGCTT | This work |
| Tick_225 | TTTGAGTGCTGGGGCGTGTT | This work |
| Tick_226 | CAGACCTTCCCAGGGGAGTG | This work |
| Tick_227 | TTCGGATGTGCCTTTTTCTT | This work |
| Tick_228 | GCTTCATTTCTTCTTGGATT | This work |
| Tick_229 | GCAAGAACCGCACTGCTCTA | This work |
| Tick_230 | AAAAATTTAATTTTTAAACT | This work |
| Tick_231 | AAGGAGGAGCGAACCGGCCC | This work |
| Tick_232 | GACGTACTCGATTACCGGCC | This work |

| | | |
|---|---|---|
| Tick_233 | GTTCGATAAAAACTCGTAGG | This work |
| Tick_234 | CTAGACGGACGGGTTGAGGC | This work |
| Tick_235 | GCAACAACGCCACCCGAAGG | This work |
| Tick_236 | TTTATGAAATTAAAAAATTT | This work |
| Tick_237 | TATCTGATACGGGTCCTATT | This work |
| Tick_238 | CCATCCAGGGGATGACGTTT | This work |
| Tick_239 | GAAGACTCGCTGCTTGGCCG | This work |
| Tick_240 | GGTTTACCTCACCTATGCCA | This work |
| Tick_241 | AAACGATGCCAAATAACATC | This work |
| Tick_242 | GGTAGGTTATGCGCCTACCA | This work |
| Tick_243 | AAAATCTTTCGCACGTTGAA | This work |
| Tick_244 | AAACCACAAACAAAGAAAAC | This work |
| Tick_245 | TCGCATCGGCGTCTCCGAGC | This work |
| Tick_246 | GGCGTTGGCGCGTTGAAACG | This work |
| Tick_247 | GCCCACCGTCGGACACCCAC | This work |
| Tick_248 | AGTCTTTCGCCCCTATACCT | This work |
| Tick_249 | ACGACATCAGTGTCGCTTGC | This work |
| Tick_250 | TCGATCGATTCTGTTTTCTT | This work |
| Tick_251 | CGATCTGAGCTCGATTTGCA | This work |
| Tick_252 | GTGGCCCCGCGTCCGCGCGT | This work |
| Tick_253 | GACGGCACGACACAACCGCG | This work |
| Tick_254 | GTTTTGACTGTGTCGGATCG | This work |
| Tick_255 | AATCACGAAGGAGGTTCCAC | This work |
| Tick_256 | TGAATTGTCGACCTCACATC | This work |
| Tick_257 | CGAAGATCAGGAATATGCGG | This work |
| Tick_258 | CGATCTCTGGCGTCTGCCGT | This work |
| Tick_259 | AAAGAAAAAACGCTTCTGGG | This work |
| Tick_260 | GACGGCATCCAAGCTTGGTC | This work |
| Tick_261 | AAGTGCACCGGGACAATCCG | This work |
| Tick_262 | ATGGTGAACTATGCCCGGGC | This work |
| Tick_263 | TGTATTTTAAAAATTTATAA | This work |
| Tick_264 | CTAATGCCCCCACGTCCACG | This work |
| Tick_265 | TCCGGAGGCCTCAGACCAGA | This work |
| Tick_266 | GAGGACTGAGCCGGTCGGGC | This work |
| Tick_267 | TCATCATGGCCCTTATGTCC | This work |
| Tick_268 | GCTGGAATTACCGCGGCTGC | This work |
| Tick_269 | TGAGACAAGCATATGACTAC | This work |
| Tick_270 | AGCCTATCCCCGGCCGCTGC | This work |
| Tick_271 | AGTATTCATCGGGGACGCGA | This work |

| | | |
|---|---|---|
| Tick_272 | TCTTTGAGCAAATGCACGAG | This work |
| Tick_273 | CACCTTGCACCCTTCTTAGA | This work |
| Tick_274 | TGATGTCGACCGAACACAGT | This work |
| Tick_275 | TTTACGGGCTCTCCTTGTGG | This work |
| Tick_276 | ATATCATTGCACTGCTTAGA | This work |
| Tick_277 | AGATCCTGTTTCTGTTCTGT | This work |
| Tick_278 | CTGCAGAGCGTTGCATTTAT | This work |
| Tick_279 | CTCCTGTAGGTTACTAAAGT | This work |
| Tick_280 | GGATAACAGCGTAATAATTT | This work |
| Tick_281 | GGTAAGCTGCGAAAAGCTTG | This work |
| Tick_282 | TTCTTTTCAACTTTCCCTCA | This work |
| Tick_283 | ACTCGTCATTTCACACCGTT | This work |
| Tick_284 | GTGCCCAAAGTGCTATGCTT | This work |
| Tick_285 | GGATATATATATATTCGTCA | This work |
| Tick_286 | GCCGTTGCTGCTCCCTTCAT | This work |
| Tick_287 | TGATACGGTGAATACGTTCT | This work |
| Mouse_1 | AGTTGGTGGAGCGATTTGTC | Dynerman et al. 2020 |
| Mouse_2 | CGTGGTCACCATGGTAGGCA | Dynerman et al. 2020 |
| Mouse_3 | GGTAAGCAGAACTGGCGCTG | Dynerman et al. 2020 |
| Mouse_4 | GTTTATGGTCGGAACTACGA | Dynerman et al. 2020 |
| Mouse_5 | GGAGGGAACCAGCTACTAGA | Dynerman et al. 2020 |
| Mouse_6 | CACAGTTATCCAAGTAGGAG | Dynerman et al. 2020 |
| Mouse_7 | TCCCCGATCCCCATCACGAA | Dynerman et al. 2020 |
| Mouse_8 | CTCCAATGGATCCTCGTTAA | Dynerman et al. 2020 |
| Mouse_9 | TTTTTCAAAGTAAACGCTTC | Dynerman et al. 2020 |
| Mouse_10 | GCAGTTGAACATGGGTCAGT | Dynerman et al. 2020 |
| Mouse_11 | TGTTATTGCTCAATCTCGGG | Dynerman et al. 2020 |
| Mouse_12 | AGACATTTGGTGTATGTGCT | Dynerman et al. 2020 |
| Mouse_13 | TCTAGAGTCACCAAAGCCGC | Dynerman et al. 2020 |
| Mouse_14 | TTCTTTTCAACTTTCCCTTA | Dynerman et al. 2020 |
| Mouse_15 | TCAGGGCTAGTTGATTCGGC | Dynerman et al. 2020 |
| Mouse_16 | GAGGCCTCGGGATCCCACCT | Dynerman et al. 2020 |
| Mouse_17 | TAGAGGTGAAATTCTTGGAC | Dynerman et al. 2020 |
| Mouse_18 | GTTTCGCTGGATAGTAGGTA | Dynerman et al. 2020 |
| Mouse_19 | TTACCGCACTGGACGCCTCG | Dynerman et al. 2020 |
| Mouse_20 | TGACCTTTTGGGTTTTAAGC | Dynerman et al. 2020 |
| Mouse_21 | CTGGCATGTTGGAACAATGT | Dynerman et al. 2020 |
| Mouse_22 | CTTTAAATGGGTAAGAAGCC | Dynerman et al. 2020 |
| Mouse_23 | GGCGAAGCCAGAGGAAACTC | Dynerman et al. 2020 |

| | | |
|---|---|---|
| Mouse_24 | CATCGCGTCAACACCCGCCG | Dynerman et al. 2020 |
| Mouse_25 | CAAAGTCTTTGGGTTCCGGG | Dynerman et al. 2020 |
| Mouse_26 | ATTCCCAAGCAACCCGACTC | Dynerman et al. 2020 |
| Mouse_27 | GGTTCTATTTTGTTGGTTTT | Dynerman et al. 2020 |
| Mouse_28 | AAAAGCTCGTAGTTGGATCT | Dynerman et al. 2020 |
| Mouse_29 | AAGGGCCAGCGAGAGCTCAC | Dynerman et al. 2020 |
| Mouse_30 | ATCCAATCGGTAGTAGCGAC | Dynerman et al. 2020 |
| Mouse_31 | CCCCGAGGGGCTCTCGCTTC | Dynerman et al. 2020 |
| Mouse_32 | AGGTGTCCTAAGGCGAGCTC | Dynerman et al. 2020 |
| Mouse_33 | AGCTTGACTCTAGTCTGGCA | Dynerman et al. 2020 |
| Mouse_34 | CGAAACCCCGACCCAGAAGC | Dynerman et al. 2020 |
| Mouse_35 | TTTTTCGTCACTACCTCCCC | Dynerman et al. 2020 |
| Mouse_36 | CGTGCGTACTTAGACATGCA | Dynerman et al. 2020 |
| Mouse_37 | GAGAACTTTGAAGGCCGAAG | Dynerman et al. 2020 |
| Mouse_38 | GGCCCAAGTCCTTCTGATCG | Dynerman et al. 2020 |
| Mouse_39 | GAAAGATCCGCCGGGACCAC | Dynerman et al. 2020 |
| Mouse_40 | ATAGCTCTTTCTCGATTCCG | Dynerman et al. 2020 |
| Mouse_41 | GCGACGTCGCTATGAACGCT | Dynerman et al. 2020 |
| Mouse_42 | GTCTGATGAGCGTCGGCATC | Dynerman et al. 2020 |
| Mouse_43 | GCAACAACACATCATCAGTA | Dynerman et al. 2020 |
| Mouse_44 | ATCAGGGTTCGATTCCGGAG | Dynerman et al. 2020 |
| Mouse_45 | ACGCTCGTGCTCCACCTCCC | Dynerman et al. 2020 |
| Mouse_46 | GCCAGAGTCTCGTTCGTTAT | Dynerman et al. 2020 |
| Mouse_47 | AAGTCAGAATCCCGCCCAGG | Dynerman et al. 2020 |
| Mouse_48 | AGCTTATGACCCGCACTTAC | Dynerman et al. 2020 |
| Mouse_49 | GGTCGAACTTGACTATCTAG | Dynerman et al. 2020 |
| Mouse_50 | GAAGTTTCCCTCAGGATAGC | Dynerman et al. 2020 |
| Mouse_51 | TGAGAGTAGTGGTATTTCAC | Dynerman et al. 2020 |
| Mouse_52 | GCGATGGCCTCCGTTGCCCT | Dynerman et al. 2020 |
| Mouse_53 | CATTAATCAAGAACGAAAGT | Dynerman et al. 2020 |
| Mouse_54 | CAGCCGACTTAGAACTGGTG | Dynerman et al. 2020 |
| Mouse_55 | GTGTAGCGCGCGTGCAGCCC | Dynerman et al. 2020 |
| Mouse_56 | CTAACGCGTGCGCGAGTCAG | Dynerman et al. 2020 |
| Mouse_57 | GCTTTTGCCCTTCTGCTCCA | Dynerman et al. 2020 |
| Mouse_58 | CTGATCTCGGAAGCTAAGCA | Dynerman et al. 2020 |
| Mouse_59 | TAAGGATTGGCTCTAAGGGC | Dynerman et al. 2020 |
| Mouse_60 | TTGTGAAGGGCAGGGCGCCC | Dynerman et al. 2020 |
| Mouse_61 | TTATCAGATCAAAACCAACC | Dynerman et al. 2020 |
| Mouse_62 | CGAATGGGTCGTCGCCGCCA | Dynerman et al. 2020 |

| | | |
|---|---|---|
| Mouse_63 | TTAATATACGCTATTGGAGC | Dynerman et al. 2020 |
| Mouse_64 | CCAGGTGGGGAGTTTGACTG | Dynerman et al. 2020 |
| Mouse_65 | CAACTTCTTAGAGGGACAAG | Dynerman et al. 2020 |
| Mouse_66 | AATGGATGGCGCTGGAGCGT | Dynerman et al. 2020 |
| Mouse_67 | TGCAAATCGGTCGTCCGACC | Dynerman et al. 2020 |
| Mouse_68 | CGGACGGGGACCCGGCTATC | Dynerman et al. 2020 |
| Mouse_69 | TGTTGGTTGATATAGACAGC | Dynerman et al. 2020 |
| Mouse_70 | ATGGAGTTTACCACCCGCTT | Dynerman et al. 2020 |
| Mouse_71 | CCGGTAAAGCGAATGATTAG | Dynerman et al. 2020 |
| Mouse_72 | CTCTTTCGAGGCCCTGTAAT | Dynerman et al. 2020 |
| Mouse_73 | GTGATTATGCTACCTTTGCA | Dynerman et al. 2020 |
| Mouse_74 | GAAACCGTTAAGAGGTAAAC | Dynerman et al. 2020 |
| Mouse_75 | GAGAGGCAAGGGGCGGGGAC | Dynerman et al. 2020 |
| Mouse_76 | AGGCACTCGCATTCCACGCC | Dynerman et al. 2020 |
| Mouse_77 | CGGGCTTGGCGGAATCAGCG | Dynerman et al. 2020 |
| Mouse_78 | TTGAGACAAGCATATGCTAC | Dynerman et al. 2020 |
| Mouse_79 | CTGCGGATATGGGTACGGCC | Dynerman et al. 2020 |
| Mouse_80 | CAAATTACCCACTCCCGACC | Dynerman et al. 2020 |
| Mouse_81 | AATAAGACGAGAAGACCCTA | Dynerman et al. 2020 |
| Mouse_82 | CTTTGCAACCATACTCCCCC | Dynerman et al. 2020 |
| Mouse_83 | CGGTTCCTCTCGTACTGAGC | Dynerman et al. 2020 |
| Mouse_84 | CTTGGCAAATGCTTTCGCTC | Dynerman et al. 2020 |
| Mouse_85 | GCACGGTGAAGAGACATGAG | Dynerman et al. 2020 |
| Mouse_86 | GCGTTATTCCCATGACCCGC | Dynerman et al. 2020 |
| Mouse_87 | GGGCGCCTTAACCCGGCGTT | Dynerman et al. 2020 |
| Mouse_88 | CAAAGCAGGCCCGAGCCGCC | Dynerman et al. 2020 |
| Mouse_89 | TAAATCAGTTATGGTTCCTT | Dynerman et al. 2020 |
| Mouse_90 | CATTGCGCCACGGCGGCTTT | Dynerman et al. 2020 |
| Mouse_91 | GTGGAGGGGTCGGGAGGAAC | Dynerman et al. 2020 |
| Mouse_92 | GCAATTATTCCCCATGAACG | Dynerman et al. 2020 |
| Mouse_93 | TTCAGTCATAATCCCACAGA | Dynerman et al. 2020 |
| Mouse_94 | ACGAATACAGACCGTGAAAG | Dynerman et al. 2020 |
| Mouse_95 | GAACTGAGGCCATGATTAAG | Dynerman et al. 2020 |
| Mouse_96 | GACTACCATCGAAAGTTGAT | Dynerman et al. 2020 |
| Borrelia_1 | TGATACGGTGAATACGTTCT | This work |
| Borrelia_2 | TCATCATGGCCCTTATGTCC | This work |
| Borrelia_3 | AGCTTGACACATTTAAAGTT | This work |
| Borrelia_4 | GTAAGCTGCGAAAAGCTTGG | This work |
| Borrelia_5 | GAGACCTTAGCTGTTGGTCT | This work |

| | | |
|---|---|---|
| Borrelia_6 | AGTGAAAGGCTAAACAAACT | This work |
| Borrelia_7 | CGTACCTCTTATAGTTTCGA | This work |
| Borrelia_8 | GCTGGGGAGACCACTCTCCA | This work |
| Borrelia_9 | GCTTCTAAGGTTAGGCTATA | This work |
| Borrelia_10 | CCTGTATATAGACCCCAAAC | This work |
| Borrelia_11 | ATAGCTAGCATCTTGCTAGC | This work |
| Borrelia_12 | AATGCTATTAAATGATGAAT | This work |
| Borrelia_13 | GGATGATCTACCTATGAGAT | This work |
| Borrelia_14 | CTTAAGCATGCAAGTCAAAC | This work |
| Borrelia_15 | GAGAGGGTGAACGGTCACAC | This work |
| Borrelia_16 | CGCAATGGGCGAAAGCCTGA | This work |
| Borrelia_17 | TCACATCTTAGCTCTCTTAA | This work |
| Borrelia_18 | GAAAGTAGGTCTTAGTGATC | This work |
| Borrelia_19 | GGCTGAACTTAAATCCATTT | This work |
| Borrelia_20 | CATGCTGGTAACAGATAACA | This work |
| Borrelia_21 | AGCGAAAGCGAGTCTTAAAA | This work |
| Borrelia_22 | GAAACCTAATTAAACAAGGG | This work |
| Borrelia_23 | GCATAAAGCAGGTCTCAGTC | This work |
| Borrelia_24 | GGGTAAAGGTCGTCTTCCCA | This work |
| Borrelia_25 | TACAGGACTATCACCTTCTT | This work |
| Borrelia_26 | GTCTTATTAGCTAGTTGGTA | This work |
| Borrelia_27 | CTAGAGTTCGGCCCTCCACT | This work |
| Borrelia_28 | CTCTTTGATTTCTTTTCCTC | This work |
| Borrelia_29 | TAAGTACGACCTAAGCAATT | This work |
| Borrelia_30 | GGTATGAGTAACGAAAAAAT | This work |
| Borrelia_31 | TTTGACTGCTTGTAAGTCTA | This work |
| Borrelia_32 | TAGGTGGGATGAAAATTCTA | This work |
| Borrelia_33 | TATGGGGTTGTAGGACGTTT | This work |
| Borrelia_34 | TTAATTTGTTAATTGATGAA | This work |
| Borrelia_35 | GTCGATGTGAACTCTTGGGA | This work |
| Borrelia_36 | GATCTTAACGAAAGTTAAGA | This work |
| Borrelia_37 | TTGTCCTAGTTTAAGCATTA | This work |
| Borrelia_38 | AAAATACCACAGCTCAACTG | This work |
| Borrelia_39 | AGCTCGTGCTGTGAGGTGTT | This work |
| Borrelia_40 | GCGCACCTCCGTTACTCTTT | This work |
| Borrelia_41 | GTTGATATCAGAAAGAATAC | This work |
| Borrelia_42 | ATTTATATAGAAGAATAATC | This work |
| Borrelia_43 | TCGCAGGTTCATTATGCAAA | This work |
| Borrelia_44 | AAATCTGTAAAGAGAAGTAT | This work |

| | | |
|---|---|---|
| Borrelia_45 | TCTGTAAGTGTAAAGGCATA | This work |
| Borrelia_46 | AAGTGTAGTCGATGGGAAAC | This work |
| Borrelia_47 | AAATGAGGAATAAGCTTTGT | This work |
| Borrelia_48 | TGATAAGTCGGAGGAAGGTG | This work |
| Borrelia_49 | ATAAATAGATCACTCGGCTT | This work |
| Borrelia_50 | TAAATACCTTCCTCCCTTAC | This work |
| Borrelia_51 | TTTTGCAGAGTTCCTTAACG | This work |
| Borrelia_52 | TGTTTCGCTTCGCTTTGTAC | This work |
| Borrelia_53 | GAAACTCGACTTCATGAAGT | This work |
| Borrelia_54 | GTTAATTTATGAATAAGCCC | This work |
| Borrelia_55 | CGACACTGCGTGAATGAAGA | This work |
| Borrelia_56 | ATTGAGCTAGGGCCTGTCAA | This work |
| Borrelia_57 | ATTTAGGTAGCATGAAATGT | This work |
| Borrelia_58 | TAAACTGGAAAGTTTGATGG | This work |
| Borrelia_59 | AATAATAGAGGCGATACCAG | This work |
| Borrelia_60 | TGGCGAACAGCCATACCCTT | This work |
| Borrelia_61 | AAATGTTCCTGAGTAGGACG | This work |
| Borrelia_62 | TGAATTTAAGCCCCAGTAAA | This work |
| Borrelia_63 | ATATGTCTAGAGTCTGATAG | This work |
| Borrelia_64 | ATTGGGCGTAAAGGGTGAGT | This work |
| Borrelia_65 | TGCTGCCTCCCGTAGGAGTC | This work |
| Borrelia_66 | ATACGCGAGGAACCTTACCA | This work |
| Borrelia_67 | TTGATCCCAAGATTACCGAA | This work |
| Borrelia_68 | GGAGGCGAAGGCGAACTTCT | This work |
| Borrelia_69 | CTTTGAAGCTATCTCGTCAG | This work |
| Borrelia_70 | GCCGTCACTTAACGAATAAA | This work |
| Borrelia_71 | CGATAAGGTTCATAGTCGAG | This work |
| Borrelia_72 | AGAAATATGAGGAGAGTGCC | This work |
| Borrelia_73 | CTCAACTCGGGTGGTGTGAC | This work |
| Borrelia_74 | TAAACACATTACCGAAGCTT | This work |
| Borrelia_75 | GTTTAGGTACGTAAACAGCC | This work |
| Borrelia_76 | ATAGTAGCTAATACCGAATA | This work |
| Borrelia_77 | AGATTAAAGCATAGAAGTGC | This work |
| Borrelia_78 | AGTTAAGCTGGAAAGTTTGA | This work |
| Borrelia_79 | GGTAGCCGTACTGGAAAGTG | This work |
| Borrelia_80 | GTTTTTGGTAAACAGTCGTT | This work |
| Borrelia_81 | GGATGTAGCAATACATTCAG | This work |
| Borrelia_82 | TTTTATTTAAAAAAATTAAC | This work |
| Borrelia_83 | TTAGTAGTGGCGAGCGAAAA | This work |

| | | |
|---|---|---|
| Borrelia_84 | ATCTACAAGCGAAGCTTTAA | This work |
| Borrelia_85 | TTACTTATCATTGCCTTGGT | This work |
| Borrelia_86 | GCATGAGGTTAACTACTGCT | This work |
| Borrelia_87 | GGCAGGGATGAGCTGTGAAT | This work |
| Borrelia_88 | ACAAGCGTTGGATATTTGAG | This work |
| Borrelia_89 | AGTCGAACAGATACGAAAGT | This work |
| Borrelia_90 | ATCCTATATATGTCAAGCCC | This work |
| Borrelia_91 | AAGGTAGCGAAATTCCTTGT | This work |
| Borrelia_92 | ACCTTAGGTTTTCGGCGAAT | This work |
| Borrelia_93 | GTTAAGCTCTTATTCGCTGA | This work |
| Borrelia_94 | AGGGTTTGGCACCTCGATGT | This work |
| Borrelia_95 | CCTCTTAACCTTCCAGCACC | This work |
| Borrelia_96 | CACACTTAACACGTTAGCTT | This work |

**Table S2:** Sequencing statistics from STAR after aligning reads to the *Ixodes scapularis* ISE6 genome.

| Sample name | Replicate | Million Seqs | % Aligned | M Aligned | % Assigned | M Assigned | % Duplicated reads |
|---|---|---|---|---|---|---|---|
| $+Bb_{mouse}$ | 1 | 11.7 | 32.40% | 4.6 | 39.40% | 2.6 | 0.8835 |
| $+Bb_{mouse}$ | 2 | 27.7 | 51.90% | 17.8 | 41.90% | 10.9 | 0.88 |
| $+Bb_{mouse}$ | 3 | 12.4 | 39.40% | 6.1 | 33.00% | 3 | 0.8555 |
| $+Bb_{lizard}$ | 1 | 18.1 | 51.30% | 11.6 | 40.70% | 7 | 0.8725 |
| $+Bb_{lizard}$ | 2 | 18.3 | 51.10% | 11.6 | 41.50% | 7.1 | 0.8715 |
| $+Bb_{lizard}$ | 3 | 19.8 | 54.00% | 13.8 | 38.60% | 8.4 | 0.8825 |
| $-Bb_{mouse}$ | 1 | 3.4 | 46.80% | 2.1 | 35.90% | 1.2 | 0.832 |
| $-Bb_{mouse}$ | 2 | 25 | 35.10% | 12.1 | 14.40% | 3.4 | 0.7575 |
| $-Bb_{mouse}$ | 3 | 6 | 41.80% | 3.2 | 36.00% | 1.8 | 0.872 |
| $-Bb_{lizard}$ | 1 | 19.4 | 52.80% | 12.9 | 40.60% | 7.8 | 0.8835 |
| $-Bb_{lizard}$ | 2 | 18.6 | 48.20% | 10.8 | 43.10% | 6.5 | 0.8645 |
| $-Bb_{lizard}$ | 3 | 6.2 | 48.20% | 3.7 | 42.00% | 2.3 | 0.857 |
| $UF_{mouse}$ | 1 | 21.4 | 48.00% | 11.9 | 37.90% | 5.9 | 0.853 |
| $UF_{mouse}$ | 2 | 20.8 | 48.00% | 11.8 | 38.90% | 6.1 | 0.819 |
| $UF_{mouse}$ | 3 | 22.1 | 47.90% | 12.4 | 38.20% | 6.1 | 0.8555 |
| $UF_{lizard}$ | 1 | 20.3 | 47.40% | 11.3 | 38.40% | 5.7 | 0.843 |
| $UF_{lizard}$ | 2 | 18.3 | 45.00% | 9.6 | 37.70% | 4.7 | 0.86 |
| $UF_{lizard}$ | 3 | 25.6 | 48.80% | 14.5 | 39.60% | 7.5 | 0.8505 |
